# Dopey-dependent regulation of extracellular vesicles maintains neuronal morphology

**DOI:** 10.1101/2024.05.07.591898

**Authors:** Seungmee Park, Nathaniel Noblett, Lauren Pitts, Antonio Colavita, Ann M. Wehman, Yishi Jin, Andrew D. Chisholm

**Affiliations:** Department of Neurobiology, School of Biological Sciences, University of California San Diego, La Jolla, CA 92093, USA; Neuroscience Program, Ottawa Hospital Research Institute, University of Ottawa, Ottawa, Ontario, Canada; Department of Biological Sciences, University of Denver, Denver, CO 80208, USA

**Keywords:** neuronal maintenance, endosomal trafficking, TAT-5 phospholipid flippase, DIP-2 lipid regulator, SAX-2/Fry, PAD-1/DOPEY

## Abstract

Mature neurons maintain their distinctive morphology for extended periods in adult life. Compared to developmental neurite outgrowth, axon guidance, and target selection, relatively little is known of mechanisms that maintain mature neuron morphology. Loss of function in *C. elegans* DIP-2, a member of the conserved lipid metabolic regulator Dip2 family, results in progressive overgrowth of neurites in adults. We find that *dip-2* mutants display specific genetic interactions with *sax-2*, the *C. elegans* ortholog of Drosophila Furry and mammalian FRY. Combined loss of DIP-2 and SAX-2 results in severe disruption of neuronal morphology maintenance accompanied by increased release of neuronal extracellular vesicles (EVs). By screening for suppressors of *dip-2 sax-2* double mutant defects we identified gain-of-function (*gf*) mutations in the conserved Dopey family protein PAD-1 and its associated phospholipid flippase TAT-5/ATP9A. In *dip-2 sax-2* double mutants carrying either *pad-1(gf)* or *tat-5(gf)* mutation, EV release is reduced and neuronal morphology across multiple neuron types is restored to largely normal. PAD-1(gf) acts cell autonomously in neurons. The domain containing *pad-1*(*gf*) is essential for PAD-1 function, and PAD-1(*gf*) protein displays increased association with the plasma membrane and inhibits EV release. Our findings uncover a novel functional network of DIP-2, SAX-2, PAD-1, and TAT-5 that maintains morphology of neurons and other types of cells, shedding light on the mechanistic basis of neurological disorders involving human orthologs of these genes.

## Introduction

Neuronal morphology is established in multiple developmental steps of initial polarization, neurite outgrowth and guidance, target selection and synaptogenesis ^1^. Throughout development and in adults, neurons undergo maturation and refinement of their architecture. While many aspects of neuronal maturation remain to be elucidated ^2^, certain themes have emerged. For example, in many organisms neurons initially form excess neurites in early development that are later pruned in an activity-dependent ^3^ or stereotyped manner ^4^. In the mature nervous system, overall neuronal morphology tends to be stable, with structural plasticity generally restricted to synapses or dendritic spines ^5^. Mechanisms that maintain mature neuron or axon morphology throughout life are of high interest, given their relevance to neurological disease or neurodegeneration. Several processes promote axon maintenance, including intrinsic pathways that ensure mitochondrial quality control or the integrity of the axonal cytoskeleton ^6^. Neuronal membrane integrity must also be maintained throughout life, and mutations in lipid metabolic genes have been linked to motor neuron disease ^7^. However, relatively little is known of the mechanisms that maintain neuronal membrane morphology.

*C. elegans* is an excellent model for studies of maintenance of neuronal morphology ^8^. *C. elegans* neurons are highly stereotyped and mostly have a simple unipolar or bipolar morphology. Genes encoding cell adhesion and Wnt-signaling molecules have been shown to affect maintenance of neuron morphology ^9,10^. Other factors also inhibit neurite sprouting in mature neurons, such as the Fry ortholog SAX-2 and its associated kinase SAX-1/NDR ^11–14^, the microtubule minus-end binding protein PTRN-1/CAMSAP ^15–17^, the planar cell polarity pathway ^18^, and the lipid regulator DIP-2/DIP2 ^19^. Mutants lacking any one of these genes display largely normal early neuronal development but subsequently show progressive overgrowth of existing axons, aberrant axonal branching, sprouting of ectopic neurites, or abnormal soma morphology. Notably, complete loss of function in each of these factors often gives rise to mild or incompletely penetrant maintenance defects, suggesting that neuronal maintenance may involve functionally redundant pathways.

Among the known neuronal maintenance pathways, some involve the cytoskeleton dynamics or membrane trafficking. For example, SAX-2/FRY and SAX-1/NDR act in a conserved pathway linked to cell morphology or polarity ^20^ that may involve regulation of Hippo signaling ^21^ or microtubule organization in dividing cells ^22^. DIP2 proteins function in neuronal morphology maintenance and axon branching in *Drosophila* ^23^ and in vertebrates ^24^. DIP2 family members are thought to regulate the activation of fatty acids and/or accumulation of diacylglycerol via their adenylate-forming domains (AFDs), also known as fatty acyl-AMP ligase-like (FAAL-like, FLD) domains ^25^. Mammalian DIP2B has been implicated in neuronal tubulin acetylation in axon outgrowth ^26^. The specific function of DIP2 family members in neuronal lipid organization or membrane architecture is not known, and the relationship of DIP2 to other morphology maintenance pathways remains poorly understood.

Here, we investigate the interaction between DIP-2 and SAX-2 in neuron morphology maintenance. We show that *dip-2(0) sax-2(0)* double mutants display highly penetrant defects in adult neuron morphology. We exploit these specific synergistic phenotypes to screen for genetic suppressors and find that gain-of-function mutations in the membrane trafficking regulator PAD-1 or its associated phospholipid flippase TAT-5 can overcome *dip-2(0) sax-2(0)* double mutant defects. Combined loss of function in DIP-2 and SAX-2 leads to altered neuronal membrane architecture and elevated release of neuronal extracellular vesicles (EVs), both of which are suppressed by gain-of-function in PAD-1 or TAT-5. Our results reveal that DIP-2 and SAX-2 pathways converge on PAD-1 and TAT-5-dependent membrane trafficking to maintain morphology in neurons and non-neuronal cells.

## Results

### DIP-2/DIP2 and SAX-2/Fry function synergistically to maintain neuron morphology

To dissect the mechanisms underlying maintenance of neuronal morphology, we focused on the lateral mechanosensory touch receptor neurons (TRNs) ALM and PLM. In wild type animals, both ALM and PLM extend a single long anterior axonal process (‘axon’) with a single ventral synaptic branch (Fig 1A,B); PLM extends a long posterior process whereas ALM extends either a short (<10 µm) or no visible posterior neurite. Loss of function in *dip-2* results in progressive defects in TRN morphology, including formation of abnormally long ALM posterior neurites, termed ‘ectopic’ neurites ^19^. Similar progressive ectopic neurite defects have been reported in *sax-2* mutants, affecting the *C. elegans* Furry/Fry ortholog ^13^.

**Figure 1.**
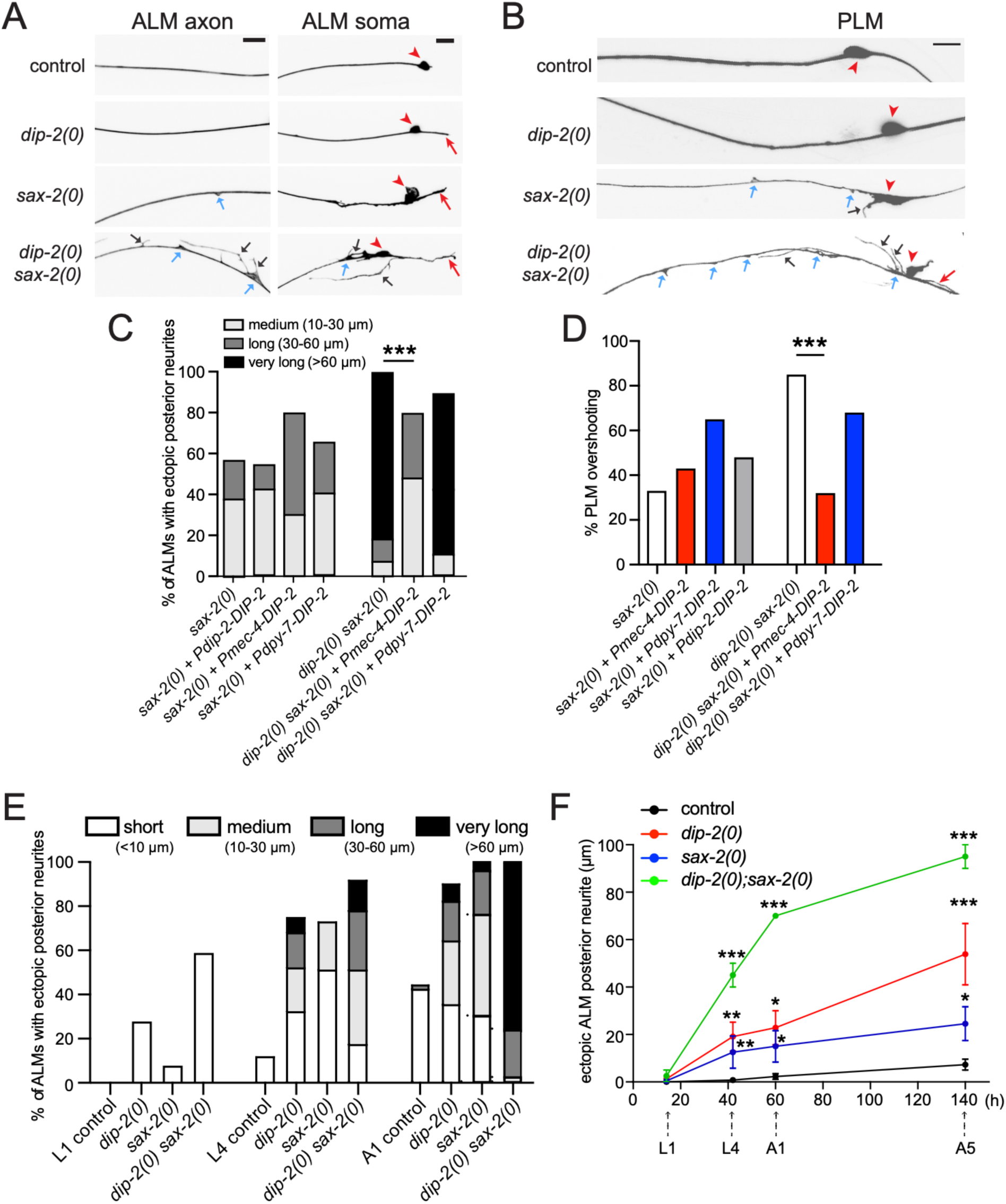
DIP-2 and SAX-2 function synergistically to maintain neuronal morphology. (A-B) Confocal images of touch receptor neurons (TRNs) labeled with P*mec-4-* GFP(*zdIs5*) in 1-day old adults of genotype indicated. Red arrowheads mark the soma of ALMs (A) and PLMs (B). In wild type, ALM extends a single anterior axonal process with a single branch in the nerve ring (not shown) and either has no posterior neurite or a very short posterior neurite (<10 µm). Any ALM posterior neurite >10 µm was scored as an ‘ectopic posterior neurite’ (red arrows). Some *sax-2(0)* mutants showed an enlarged and misshapen soma and small blebs (blue arrows) along anterior neurites. *dip-2(0) sax-2(0)* double mutants exhibited multiple ectopic neurites (black arrows) in addition to the ectopic posterior neurite, as well as large blebs (blue arrows) from the anterior axon and soma. PLM (B) showed a similar range of defects in mutants indicated. Scale = 10 µm. (C,D) Overexpression of DIP-2 did not rescue *sax-2(0)* single mutant neuronal morphology phenotypes (left columns). Overexpression of DIP-2 in TRNs but not in epidermis can partly suppress ALM ectopic posterior neurites and PLM overshooting phenotypes of *dip-2(0) sax-2(0)* double mutants to the level of *sax-2(0)* single mutants (right columns). Ectopic posterior neurites were further classified by length. Neuron morphology scored in A1 adult (N = 18-40). Statistics, Freeman-Halton extension of Fisher’s exact test. (E) Score of ALM ectopic posterior neurites in L1, L4 and 1-day old adult animals of genotype indicated, classified based on neurite length. N = 23-58. (F) Individual animals were repeatedly imaged over multiple time points to measure the length of the longest ectopic neurite in L1, L4, day 1 adult, and day 5 adult stages (N = 2-10 per time point). x-axis, hours after hatching. Statistics, one-way ANOVA. *(P<0.05), **(P<0.01), ***(P<0.001).

In a forward genetic screen for new mutants exhibiting progressive defects in TRN morphology (see Methods) we recovered two new alleles of *dip-2*, three alleles of *sax-2*, three alleles of *ptrn-1,* two alleles of the TRN-specific ß-tubulin *mec-7*, and a single allele of the MT binding protein *coel-1* (Fig S1A). While limited to viable mutants, our screen suggests that a restricted set of genes may have specific roles in TRN morphology maintenance.

To understand the relationship between these factors, we carried out genetic double mutant analysis and found a strong synergistic interaction between null (0) mutants of *dip-2* and *sax-2* (Fig 1A,B; Fig S1B). About 60% of *dip-2(0)* single mutant adults displayed an ectopic ALM posterior neurite ^19^ (red arrow, Fig 1A). About 60% of *sax-2(0)* single mutant adults also displayed ectopic ALM posterior neurites as well as small blebs or sprouts along the ALM axon (blue arrow, Fig 1A) and enlarged and deformed soma (red arrowhead, Fig 1A). In *dip-2(0) sax-2(0)* double mutants, 100% of adults displayed both an extremely elongated posterior ALM neurite and excessive sprouting or protrusions from the axon and soma (Fig 1A,E). PLM morphology in *dip-2(0) sax-2(0)* double mutants also displayed enhanced defects such that >80% of *dip-2(0) sax-2(0)* double mutants displayed PLM axon overshooting (Fig 1B,D, Table 1), defined as the anterior tip of PLM extending anterior to the ALM soma ^13^, compared to <5% of *dip-2(0)* mutants and ∼40% of *sax-2(0)* mutants (Fig 1D, Table 1). We observed similar synergistic defects using multiple loss-of-function mutations of *dip-2* and *sax-2*, including a newly generated deletion (*ju1815*) of *sax-2*, and using two TRN markers *zdIs5* and *muIs32* (Table 1, Fig S1C; see Methods). In *dip-2(0) sax-2(0)* double mutants, other types of neurons, such as GABAergic motor neurons (Fig S1D, F) and ciliated sensory neurons (Fig 6B), displayed similarly increased morphological disruption. Thus, DIP-2 and SAX-2 have overlapping functions in regulating morphology of multiple neuron types.

**Table 1.**
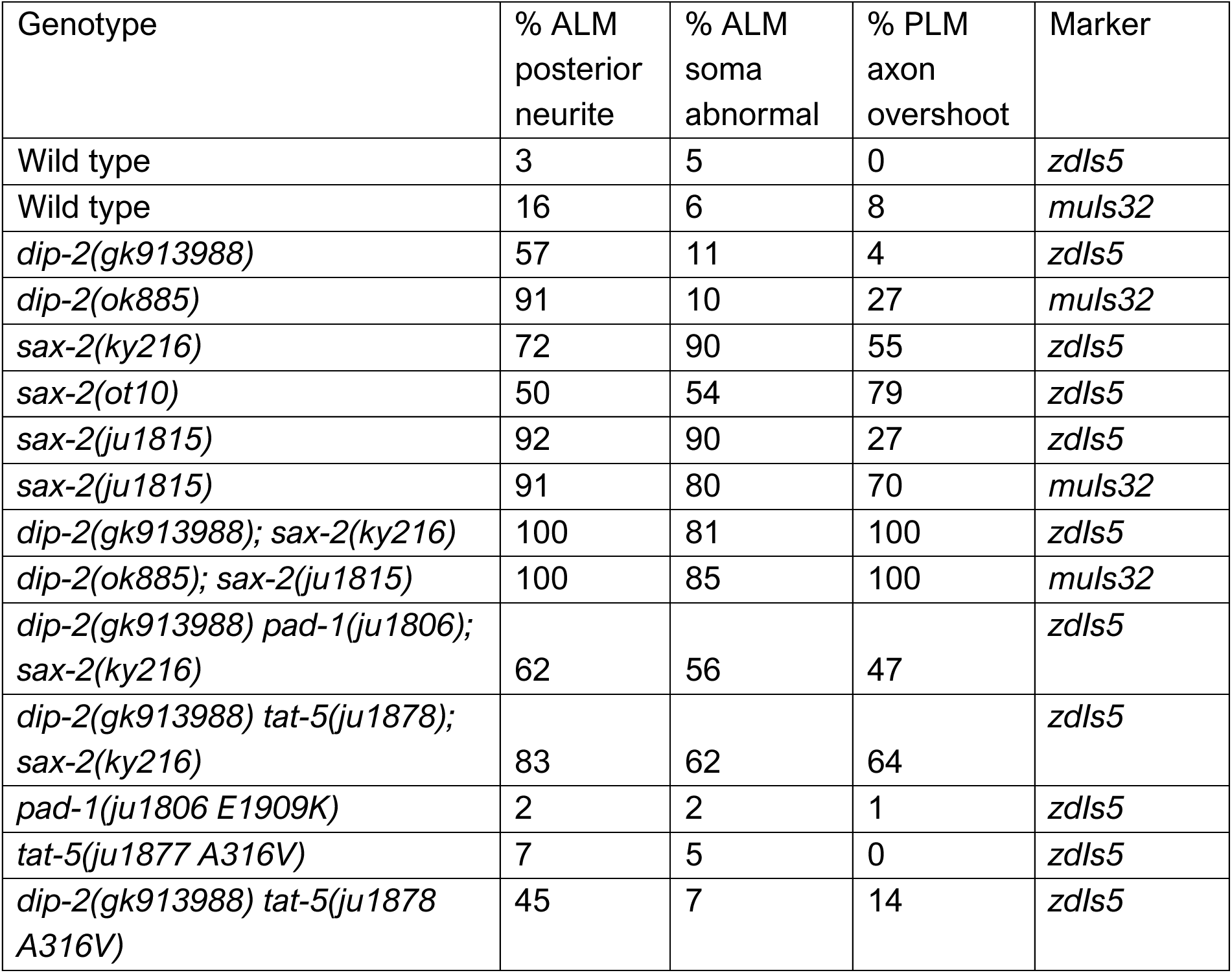
Neuronal morphology of key genotypes. See Methods for definitions of neuronal morphology defects. N = 25-65 per genotype

To explore the basis of interaction between DIP-2 and SAX-2, we overexpressed DIP-2 in *sax-2(0)* single mutants. Transgenic expression of DIP-2 under the *dip-2* promoter had been previously shown to rescue the TRN axon phenotype of *dip-2(0)* single mutants ^19^, but did not affect TRN phenotypes in *sax-2(0)* (Fig 1C,D). Expression of DIP-2(+) cDNA using the *mec-4* touch neuron-specific promoter in *dip-2(0) sax-2(0)* double mutants rescued ALM posterior neurite overgrowth and PLM axon overshooting to the level of *sax-2(0)* single mutants and did not rescue ALM soma shape defects specific to *sax-2(0)*. Conversely, expression of DIP-2 using the *dpy-7* epidermal promoter did not rescue neuronal morphology defects in *dip-2(0) sax-2(0)* (Fig 1C,D). These results are consistent with our previous findings that DIP-2 represses ectopic neurite outgrowth in a cell-autonomous manner ^19^, and suggest that mechanistically distinct regulation of a common process mediated by DIP-2 and SAX-2 restrains ectopic neurite outgrowth.

Both DIP-2 and SAX-2 function in postembryonic development to maintain neuronal morphology ^12,13,19^. We next determined the time course of ALM neuronal morphology defects in single and double mutants. In the early larval (L1) stage, single short ALM posterior neurites were observed in 30% of *dip-2(0)*, 10% of *sax-2(0)* single mutants and 60% of *dip-2(0) sax-2(0)* double mutants (Fig 1E). At the late larval (L4) stage, medium to very long ectopic neurites were observed in ∼80% *dip-2(0) sax-2(0)* double mutants; by day 1 adult, almost 100% of double mutants displayed posterior neurites > 30 µm (Fig 1E). In addition, 60% of double mutant adults displayed multiple ectopic neurites from the ALM soma (Fig 1A, Fig S1E). To track TRN morphological changes, we also followed ALM morphology in individual animals over time and found synergistic defects in *dip-2(0) sax-2(0)* double mutants that progressively increased from L4 to day 5 adults (Fig 1F), suggesting DIP-2 and SAX-2 function continuously in mature neurons.

In addition to the above defects in neuronal morphology, *dip-2(0) sax-2(0)* adults displayed shortened body length (Fig 2A and S2A,B) and reduced fertility (Table 2), compared to single mutants. Expression of DIP-2 using the *dip-2* promoter rescued body length defects of *dip-2(0) sax-2(0)* double mutants (Fig S2C). However, expression of DIP-2 under the control of either a pan-neuronal or an epidermal promoter did not rescue the body morphology phenotype (Fig S2C), suggesting this phenotype may arise from defects in multiple tissues.

**Figure 2.**
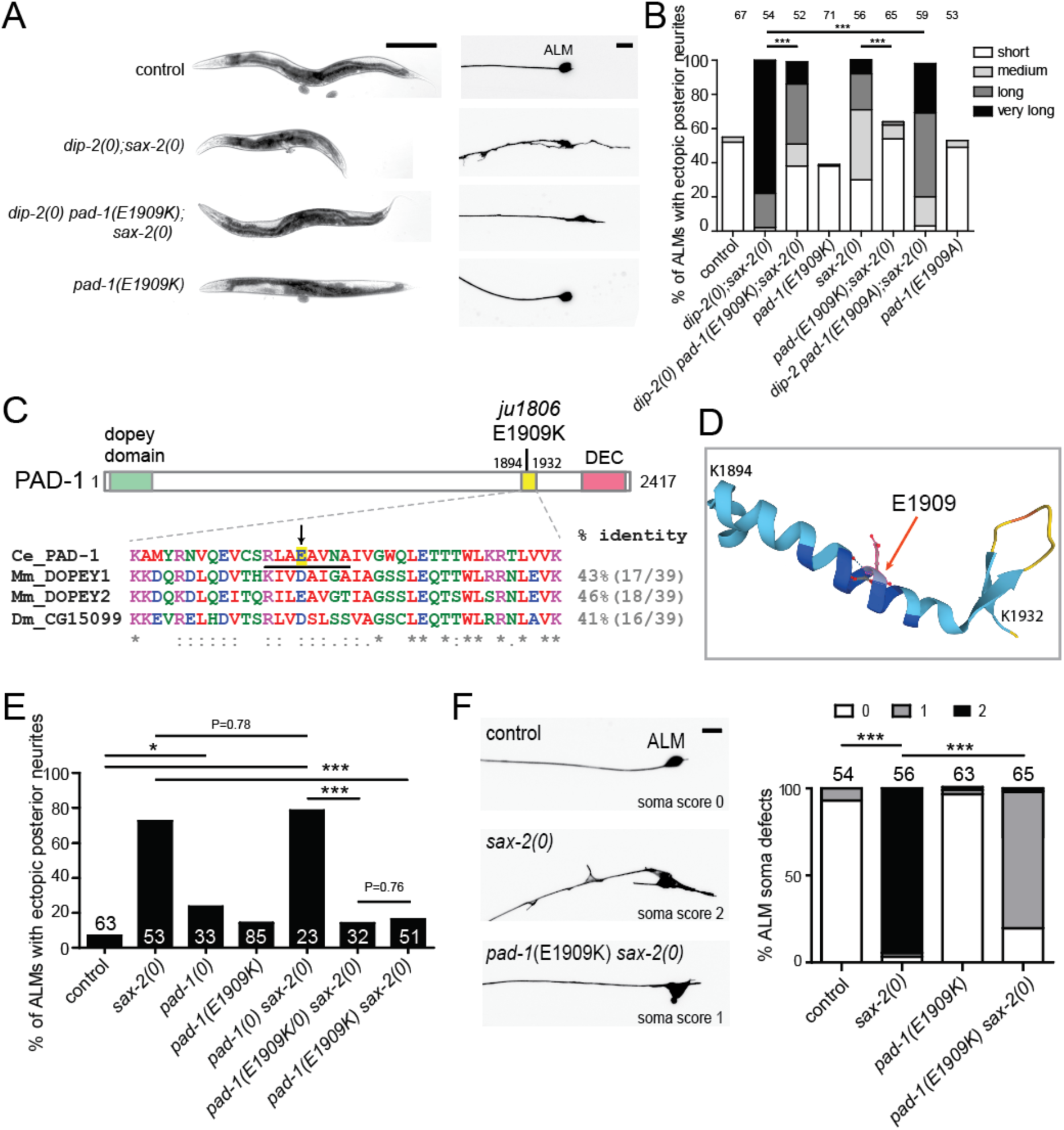
The PAD-1(E1909K) gain-of-function mutation suppresses *dip-2(0) sax-2(0)* double mutant phenotypes. (A) Bright field images of 1-day old adults (left) and confocal images of ALM neurons (right) in animals of genotype indicated. *pad-1(ju1806 E1909K)* suppressed both body length and neuronal morphology defects in *dip-2(0) sax-2(0)* double mutants. Scale = 100 µm for body images and 10 µm for neuronal images. (B) Quantitation of ALM posterior neurite phenotypes in animals of genotype indicated. *pad-1*(*E1909K*) suppresses the sprouting of long posterior neurite in *dip-2(0) sax-2(0)* double and *sax-2(0)* single mutants. *pad-1(E1909A)* reduced the length of ALM posterior neurites. (C) PAD-1 protein structure illustration with the sequence alignment below showing the region affected by *pad-1(ju1806 E1909K)*. Color-coded sequence alignment of PAD-1 residues (aa1894-1932) and the corresponding residues in the DOPEY orthologs from *Mus musculus* (UniProt Q8BL99 and Q3UHQ6) and *Drosophila melanogaster* (A1ZBE8) was obtained using Clustal Omega. % identity to PAD-1 is indicated. Mouse and human DOPEY1 are identical in this region. Arrow points to E1909, and the black underline points to 8 residues deleted in *pad-1(ju1948).* DEC corresponds to the ‘Dopey Extreme C-terminus’ implicated in targeting of Dopey1 to the Golgi ^30^. (D) Alphafold (https://alphafold.ebi.ac.uk) prediction of structure of the region spanning aa 1894-1932 in PAD-1; E1909 is indicated with red arrow. (E) Genetic interaction between *pad-1* gain and loss-of-function mutation with *sax-2(0)*. ALM ectopic neurites scored using P*mec-7*-GFP(*muIs32*) marker. *pad-1(E1909K)* displayed dominant suppression of ALM ectopic neurite outgrowth. *pad-1(0)* caused partially penetrant ectopic ALM neurites and did not enhance *sax-2(0)*. (F) *pad-1(E1909K)* suppressed soma shape defects of *sax-2(0)* single mutants. ALM soma shape defects were categorized as “0” (normal: ovoid soma shape with uniform GFP expression); “1” (soma slightly altered from the ovoid shape but even GFP expression) or “2” (soma significantly misshapen and/or uneven GFP expression). Scale = 10 µm. Statistics (D,E): Fisher’s exact test. ***(P < 0.001), **(P<0.01), *(P<0.05).

**Table 2.**
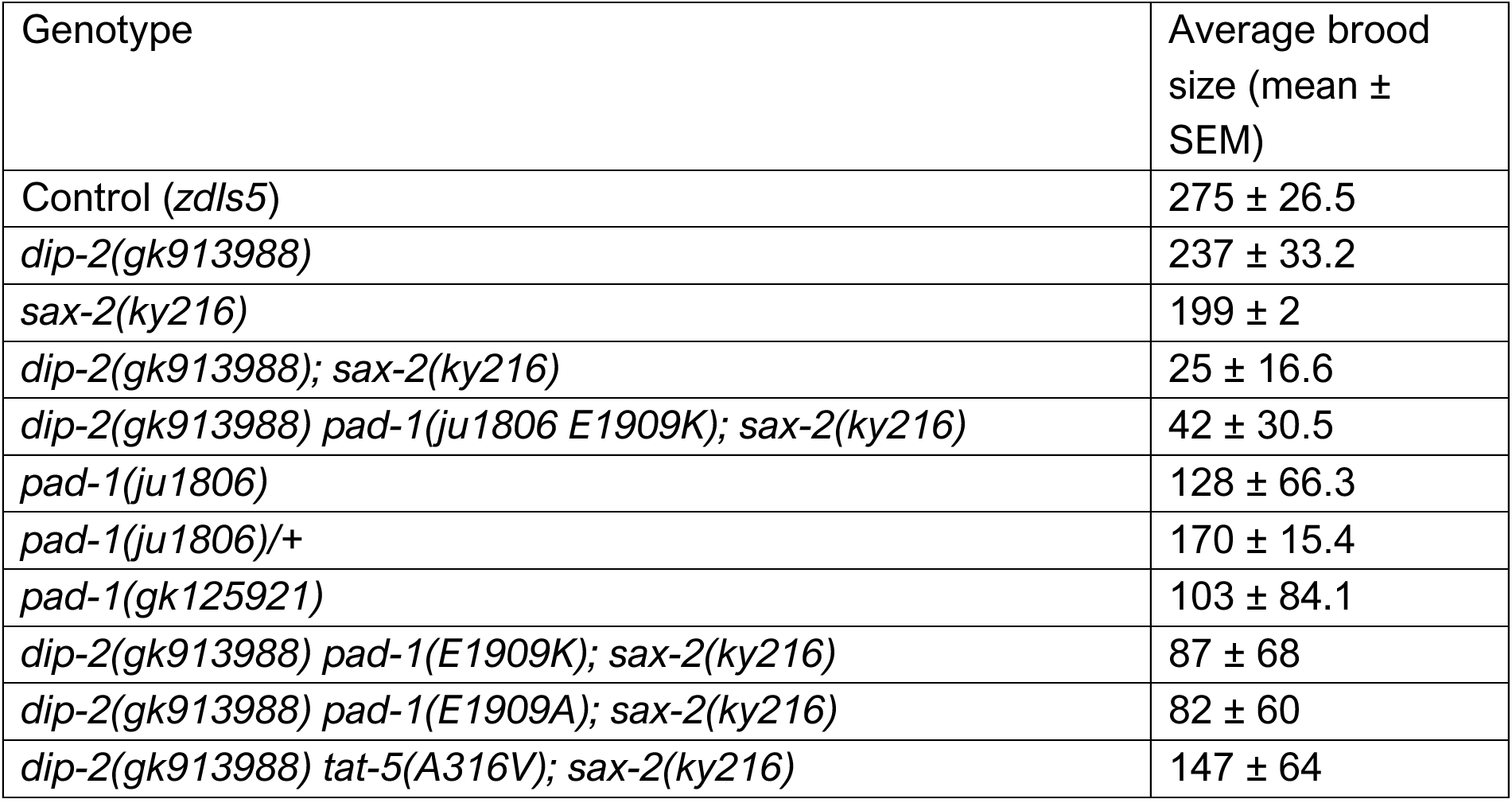
Brood size of key genotypes. N = 2-14 complete broods per genotype.

### A gain-of-function mutation in *pad-1*/Dopey suppresses *dip-2 sax-2* phenotypes

To gain insights into how DIP-2 and SAX-2 might coordinately regulate neuronal morphology, we performed a forward genetic suppressor screen (see Methods). We first screened for suppression of the shortened body length of *dip-2(0) sax-2(0)* double mutants and then examined candidate suppressors for restoration of neuronal morphology. Combining genetic mapping and whole genome sequencing, we identified one strong suppressor, *ju1806*, that ameliorated *dip-2(0) sax-2(0)* synthetic body morphology and neuronal morphology phenotypes (Fig 2A,B). This suppressor contains a G to A nucleotide transition in the *C. elegans* Dopey ortholog *pad-1*. Dopey proteins are conserved regulators of membrane trafficking, originally identified from their role in fungal cell morphology ^27^ (reviewed in ^28^). The *pad-1(ju1806)* mutation results in a missense alteration of glutamic acid at position 1909 to lysine (E1909K) (Fig 2C). GABAergic motor neuron morphology defects were also significantly suppressed in *dip-2(0) sax-2(0) pad-1(E1909K)* (Fig S1D,F). These data show that the missense alteration *pad-1(E1909K)* can restore neuronal morphology to largely normal in *dip-2(0) sax-2(0)* double mutants.

PAD-1 is essential for embryonic development, and a *pad-1(wur02)* null mutation results in recessive adult sterility or maternal-effect lethality ^29^. *pad-1(E1909K)* single mutants displayed largely normal neuronal morphology (Fig 2A, Fig S1F), slightly reduced body length (Fig S3A) and significantly smaller brood size (Table 2). Three lines of evidence suggest that *pad-1(E1909K)* causes a gain-of-function (*gf*). First, *pad-1(0)* mutant homozygotes derived from *pad-1(0)/+* heterozygous mothers displayed mild ALM ectopic neurite outgrowth (Fig 2E). *pad-1(0)* homozygotes did not suppress the ectopic neurite phenotype of *sax-2(0)* mutants but showed additive phenotypes (Fig 2E). Second, *pad-1(E1909K)/pad-1(0)* trans-heterozygotes suppressed the ALM ectopic neurite phenotype of *sax-2(0)* to the same degree as *pad-1(E1909K)* homozygotes (Fig 2E). Third, *pad-1(E1909K)* but not *pad-1(0)* behaved as a semi-dominant suppressor of *dip-2(0) sax-2(0)* double mutant phenotypes (Fig S3B). Taken together, our data suggest that PAD-1 represses ectopic neurite outgrowth and that *pad-1(E1909K)* is a gain-of-function mutation.

We also analyzed pair-wise double mutants of *pad-1(E1909K)* with *dip-2(0)* or *sax-2(0). pad-1(E1909K)* strongly suppressed the neuronal morphology defects of *sax-2(0)* single mutants, such that <20% of ALM neurons displayed ectopic posterior neurites and the majority of ALMs exhibited significantly improved soma shape (Fig 2E, F). *pad-1(E1909K)* also significantly suppressed ALM posterior neurites of *dip-2(0)* single mutants, albeit to a lesser degree (Fig 4D). These data suggest that PAD-1 may act downstream of both DIP-2 and SAX-2.

### A conserved region containing E1909 is critical for PAD-1 function

Dopey family proteins are defined by a well-conserved N-terminal Dopey domain and a C-terminal domain that is less conserved in primary sequence (Fig 2C). The N- and C-terminal regions of mammalian Dopey1 have been implicated in kinesin-1 binding and membrane association, respectively ^30^. The region containing E1909 is conserved between *C. elegans* and mammals and predicted to form an alpha-helix, but otherwise largely uncharacterized (Fig 2C,D). To test the functional importance of the region surrounding E1909, we made an in-frame deletion *pad-1(ju1948)* that removed the 8 amino acids S1905-N1912 (underlined in Fig 2C). *pad-1(ju1948)* homozygous animals lived to adults, and displayed sterility and maternal effect lethality, with 85% laying dead embryos and 15% producing no embryos (n = 20), similar to *pad-1(0)* mutants. Thus, the region affected by E1909K is essential for PAD-1 function. As the E1909K alteration causes a charge reversal, we made another genome editing to change E1909 to Ala. We found that *pad-1(E1909A)* partly suppressed the neuronal phenotypes of *dip-2(0) sax-2(0)* double mutants such that ectopic ALM posterior neurites were shorter (Fig 2B, Table 2). *pad-1(E1909A)* also suppressed the *dip-2(0) sax-2(0)* brood size defect to the same degree as *pad-1(E1909K)* (Table 2), but did not significantly suppress the body length phenotype of *dip-2(0) sax-2(0)* (Fig S3A). This result suggests that glutamic acid at the residue 1909 is crucial for normal PAD-1 function.

### SAX-2, PAD-1, and DIP-2 localize to distinct subcellular compartments

To test how DIP-2, SAX-2 and PAD-1 might interact, we examined their expression using functional fluorescent protein knock-in (KI) tagging of the endogenous loci (see Methods). We previously reported that GFP::DIP-2 KI showed diffuse localization in neurons ^19^. We generated N-terminal and C-terminal GFP KIs of SAX-2 (together denoted SAX-2::GFP) and found they were similarly expressed in multiple tissues including the germ line and the nervous system (Fig S4A). In neurons, SAX-2::GFP displayed discrete puncta in the soma (Fig 3A; Fig S4B-E) and was occasionally detected in axons. In ALM, we observed SAX-2::GFP localized to small (< 1 µm diameter) puncta clustered at the periphery of the soma (Fig 3A and Fig S4B). In other neuron types, such as the chemosensory neuron AWB, SAX-2::GFP formed fewer larger puncta in the soma (Fig S4C). We sought to identify the SAX-2::GFP-positive compartments in neurons by co-localization analysis with markers for early endosomes (RAB-5), late endosomes (RAB-7), the medial Golgi (AMAN-2), and the *trans*-Golgi (RUND-1) ^31^ (Fig S4D-G). SAX-2::GFP did not show co-localization with any of the above markers, suggesting SAX-2 puncta in neurons are unlikely to correspond to endosomes or the Golgi.

**Figure 3.**
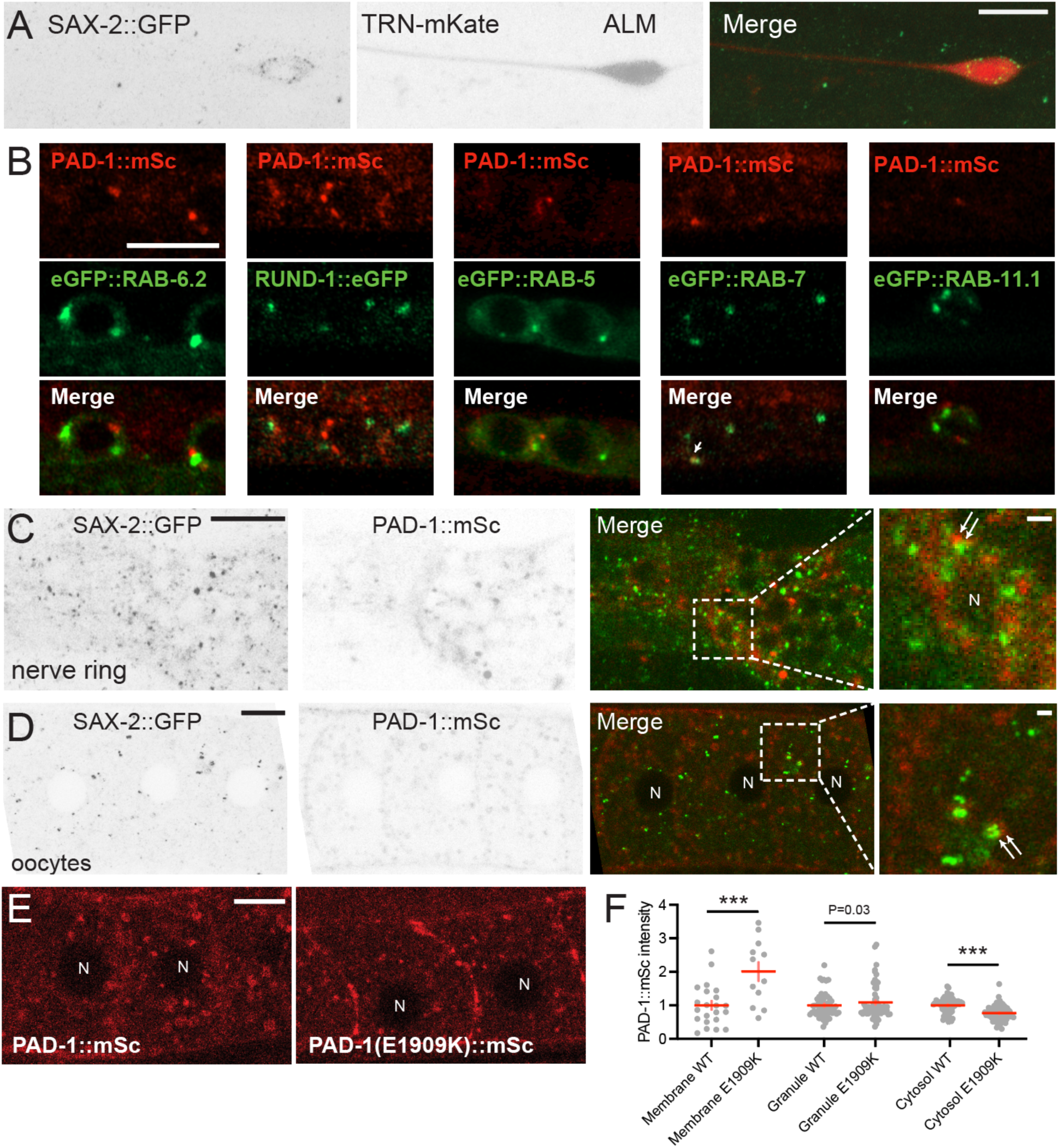
Localization of SAX-2, PAD-1(WT), and PAD-1(E1909K) (A) SAX-2::GFP(*ju1831*) in ALM neuron labelled with mKate (*juSi329*). Small SAX-2::GFP puncta were visible in the periphery of ALM soma and in surrounding epidermis. Scale, 10 µm (B) Airyscan images of PAD-1::mSc KI partially co-localized with the late endosomal marker RAB-7 in motor neurons. PAD-1 mSc KI did not co-localize with other endosomal markers RAB-5 and RAB-11.1 or Golgi markers eGFP::RAB-6.2 and RUND-1::eGFP. Scale = 5 µm. (C) Images represent the anterior ganglion near the nerve ring where N-terminal GFP::SAX-2 and PAD-1::mSc puncta are present in neuronal soma but do not co-localize. The inset image is single confocal slice of a soma (nucleus indicated by N) showing that PAD-1 and SAX-2 puncta are adjacent (white arrows) but not overlapping. Scale = 10 µm. The scale in the magnified image is 1 µm. (D) Images of oocytes showing a lack of co-localization of GFP::SAX-2 and PAD-1::mSc, with both types of puncta occasionally adjacent (white arrows). Scale = 10 µm. The scale in the magnified image is 1 µm. (E) Images of PAD-1(E1909K)::mSc in oocytes showing significantly increased association with the plasma membrane, compared to PAD-1(+)::mSc. Scale = 10 µm. (F) Quantitation of PAD-1::mSc or PAD-1(E1909K)::mSc signal intensity at the plasma membrane, cortical granules (recycling endosomes), and cytosol in oocytes. Intensities are normalized to the WT PAD-1::mSc control in each location. Statistics: t-test. ***(P < 0.001). N in white indicates oocyte nucleus.

An N-terminal GFP::PAD-1 KI is expressed in multiple tissues including neurons ^29^ and causes a partial loss of PAD-1 function ^32^. We therefore generated new KIs tagged at the C-terminus with mScarlet-I (PAD-1::mSc) or mNeonGreen (PAD-1::mNG). Both showed similarly broad expression and localized mostly to cytoplasmic puncta (Fig 3B-D; Fig S5C, S6C). Using reporters for various intracellular organelles, we observed that PAD-1 partly co-localized with a subset of RAB-7-positive puncta marking late endosomes, but not with the other endosomal markers (RAB-5 and RAB-11.1) nor the *trans*-Golgi markers (RUND-1::GFP and eGFP::RAB-6.2) (Fig 3B). This co-localization analysis suggests PAD-1 may localize to late endosomes in neurons.

We next defined the spatial relationship of SAX-2 and PAD-1. In neurons, PAD-1::mSc puncta were largely non-overlapping with SAX-2::GFP puncta, although the two types of puncta were occasionally adjacent (Fig 3C). As SAX-2 and PAD-1 were expressed at relatively low levels in neurons, we turned to oocytes where both SAX-2::GFP and PAD-1::mSc showed a punctate pattern (Fig 3D). SAX-2::GFP puncta were more uniform in size throughout the cytoplasm, often in pairs or small clusters (Fig S4A). PAD-1::mSc puncta varied in size from small (< 1 µm diameter) puncta to larger vesicles, in addition to faint expression close to the plasma membrane (Fig 3D). A subset of PAD-1+ puncta co-localized with RAB-7::mNG and RAB-11.1::GFP and also with CAV-1-labeled cortical granules (Fig S4H-J). As in neurons, PAD-1::mSc puncta in oocytes did not overlap with SAX-2::GFP puncta but were occasionally adjacent (Fig 3D). Taken together, SAX-2 and PAD-1 appear to localize to distinct subcellular compartments in neurons and oocytes.

We next addressed whether these proteins affected each other’s localization. In *dip-2(0)* mutants, neuronal SAX-2::GFP punctate distribution resembled that of the wild type (Fig S5A). Conversely, the diffused pattern of GFP::DIP-2 in the soma of TRNs was unchanged in *sax-2(0)* mutants (Fig S5B). PAD-1::mNG expression was not overtly altered in *sax-2(0)* mutants either in oocytes (Fig S5C) or in other tissues. These results suggest that DIP-2, SAX-2, and PAD-1 do not regulate each other’s localization.

### PAD-1(E1909K) acts cell-autonomously and affects PAD-1 membrane localization

Both DIP-2 and SAX-2 act cell-autonomously in neurons ^13,19^. We next tested whether PAD-1 suppressor activity is also cell-autonomous. To address this question, we initially attempted to rescue suppression of the phenotype of *sax-2(0)* by expressing full-length *pad-1* cDNA, but were not able to obtain expression, possibly due to the large size of *pad-1* cDNA and/or low translational efficiency. We then transgenically expressed *pad-1* sense and anti-sense cDNA fragments under the control of the *mec-4* TRN promoter. The suppression of morphological defects by *pad-1(E1909K)* was significantly rescued (Fig S3C), supporting the conclusion that the E1909K mutant form of PAD-1 acts in neurons.

We next examined how E1909K mutation affected PAD-1 expression, by editing this mutation in the PAD-1 KI backgrounds (see Methods). In oocytes, PAD-1(E1909K) KIs showed significantly increased association with the plasma membrane (Fig 3E,F; Fig S6A,B). PAD-1(E1909K)::mSc was not uniform along the plasma membrane but was localized to patches (Fig 3E); similar patchiness was observed with GFP::PAD-1(E1909K) (Fig S6A) and PAD-1(E1909K)::mNG (Fig S6E). While the fluorescence intensity of PAD-1(E1909K) KIs at intracellular puncta or vesicles was similar to that of the wild type PAD-1 KIs, the fluorescence intensity of cytosolic PAD-1(E1909K) KI was significantly reduced (Fig 3F; Fig S6B). This suggests that E1909K may shift PAD-1 from a cytosolic pool to a plasma membrane associated pool. In the nervous system, PAD-1(E1909K)::mNG did not show more membrane enrichment in the nerve ring than WT PAD-1::mNG (Fig S6C). We also did not detect significant differences in size and intensity between PAD-1::mNG and PAD-1(E1909K)::mNG puncta in ALM neurons, possibly due to the small size of the soma (Fig S6D). We further examined how PAD-1(E1909K) might affect the expression of GFP::DIP-2 and SAX-2::GFP, and found that *pad-1(E1909K)* did not alter the protein expression level compared to wild type (Fig S5D,E). Conversely, we did not detect significant changes in the fluorescence levels of PAD-1(E1909K)::mNG in *dip-2(0) sax-2(0)* double mutants (Fig S6E). This analysis suggests that the altered subcellular localization of PAD-1(E1909K) may enable its ability to bypass the cellular defects of *dip-2(0) sax-2(0)*.

### A gain-of-function mutation in flippase TAT-5 suppresses *dip-2 sax-2* phenotypes

*C. elegans* PAD-1 is thought to promote the activity of TAT-5, a conserved P4-ATPase flippase that maintains phosphatidylethanolamine (PE) asymmetry in the plasma membrane ^29,33^. Among additional suppressors from our screen, we found that the *ju1805* suppressor contained a missense mutation in *tat-5* (Fig 4A). The *ju1805* mutation alters a conserved alanine to valine (A316V) in the ATPase actuator (A) domain, which serves as an intramolecular phosphatase (Fig 4A) ^34^. As an independent verification of this amino acid change, we generated the same A316V alteration (denoted *ju1878*) in a *dip-2(0) sax-2(0)* double mutant background and observed a similar degree of suppression (Fig S7A). The suppression of *dip-2(0) sax-2(0)* TRN defects by *tat-5(A316V)* was overall weaker than that by *pad-1(E1909K)* (Table 1). We further generated *tat-5(A316W)* by editing and did not observe suppression of *dip-2(0) sax-2(0)* body length defects, supporting the specific effect of *tat-5(A316V)*.

**Figure 4.**
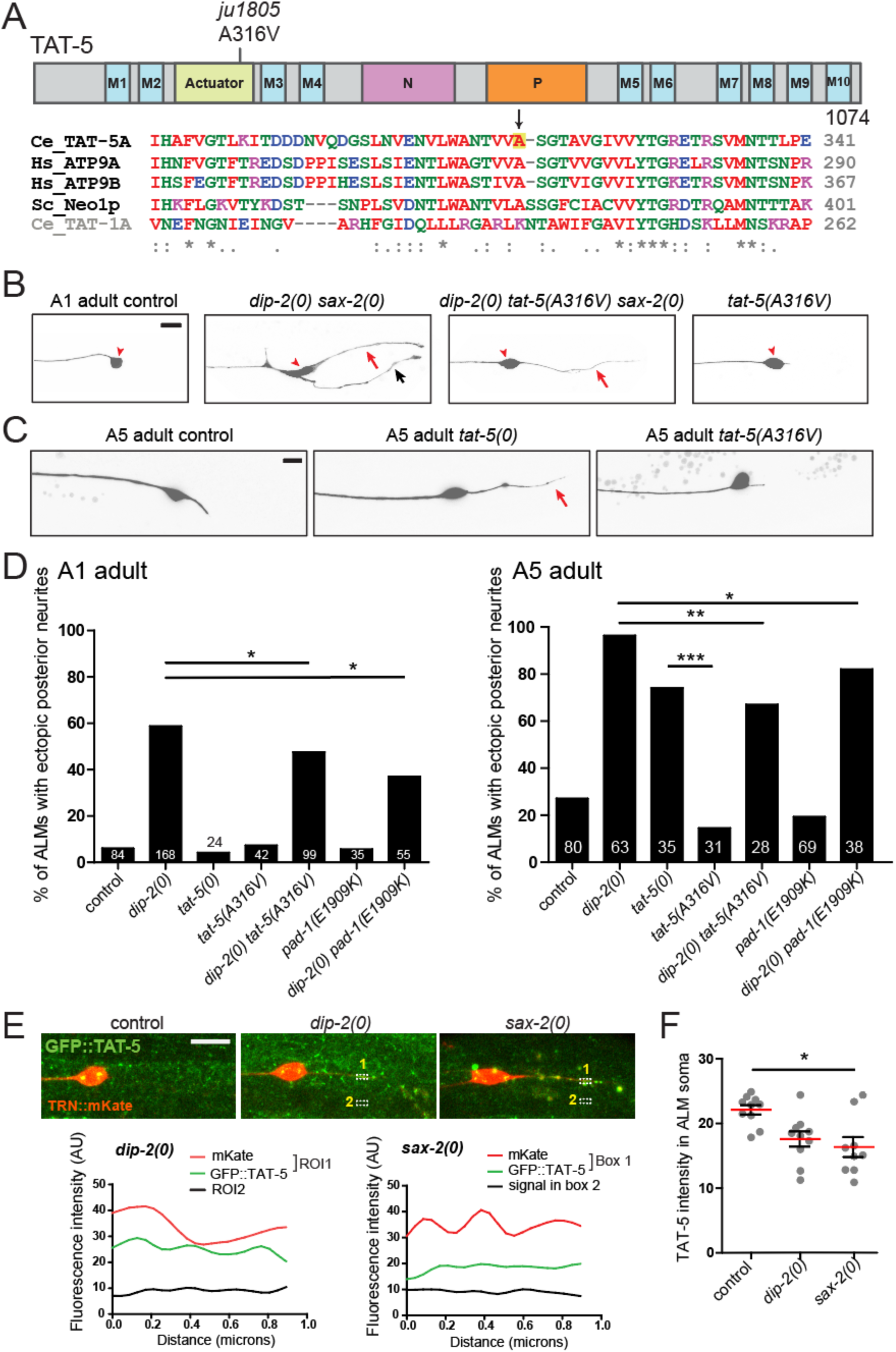
The *TAT-5(A316V)* gain-of-function mutant suppresses *dip-2(0) sax-2(0)* double mutant phenotypes. (A) Cartoon of TAT-5 isoform A. Blue M1-M10 represent the 10 transmembrane segments. The cytosolic loop between segments M2 and M3 contains part of the actuator (A) domain (yellow). The cytosolic region between M4 and M5 contains the nucleotide-binding (N) domain (pink) and the phosphorylation (P) domain (orange). Sequence alignment below shows the region encompassing A316 of TAT-5 isoform A and the corresponding region in other essential P4-ATPases of subclass 2: yeast Neo1p (QHB09325) and human ATP9A (O75110) and ATP9B (AAI25220) ^78^. A316 is conserved between TAT-5 and these orthologs, but not in TAT-1 (subclass 1b). (B) Images show suppression of ALM morphology defects in 1-day old *dip-2(0) sax-2(0)* mutant adults by *tat-5(A316V)*, including ectopic posterior neurites (red arrow), additional ectopic neurites (black arrow) and soma morphology defects (red arrowhead). *tat-5(A316V)* single mutants displayed normal ALM morphology. Scale = 10 µm. (C) Images of ALM in 5-day old control, *tat-5(0)* and *tat-5(A316V). tat-5(0)* but not *tat-5(A316V)* displayed a long posterior neurite from the ALM soma. (D) Quantitation of ALM ectopic neurite defects in aged animals of genotype indicated. Both *tat-5(A316V, ju1805) and pad-1(E1909K, ju1806)* partly suppressed *dip-2(0)* defects at 1-day old (A1, left) and 5-day old adult (A5, right) stages. ALMs in 5-day old *tat-5(A316V)* adults displayed short posterior neurites, whereas ALMs in age-matched *tat-5(0)* exhibited long posterior neurites. Statistics: Fisher’s exact test. ***(P < 0.001), **(P<0.01), *(P<0.05). (E) Airyscan images show ALM neurons labelled with *juSi329* [P*mec-4*::mKate] and expressing *wur36* [GFP::TAT-5B/D KI]. Both *dip-2(0)* and *sax-2(0)* single mutants display GFP::TAT-5 in ectopic posterior neurites. Scale = 10 µm. The fluorescent signals of GFP::TAT-5 in the “1” dashed boxes within mKate+ ectopic neurites were line-scanned and plotted to show co-localization of TAT-5 and mKate. Signals in the dashed boxes labelled “2” serve as background. (F) Quantitation of mean intensity of GFP::TAT-5 (Arbitrary Units, AU) in the ALM soma. Kruskal-Wallis test with Dunn’s post test. * (P<0.05).

*tat-5(A316V)* single mutants were viable and fertile, and displayed normal TRN morphology (Fig 4B). Like *pad-1(0)*, *tat-5(0)* mutants display sterility and maternal-effect lethality ^33^. *tat-5(A316V)/tat-5(0)* trans-heterozygous animals were healthy and produced viable progeny. To further determine whether *tat-5(A316V)* was partial loss-of-function or gain-of-function, we compared TRN morphology in *tat-5(A316V)* and *tat-5(0)*. At 1-day old adult stage, TRN morphology was normal in both mutants (Fig 4D). At 5-day old adult stage, >70% of ALM in *tat-5(0)* displayed ectopic posterior neurites, whereas only ∼10% of *tat-5(A316V)* showed such defects (Fig 4D). These results support a conclusion that TAT-5(A316V) is a gain-of-function mutation. Additionally, *tat-5(A316V)* significantly suppressed the TRN ectopic neurite defect of *dip-2(0)* single mutants both in day 1 and day 5 adults (Fig 4D) but had less effect on TRN phenotypes of *sax-2(0)* single mutants. Thus, *tat-5(A316V)* may preferentially ameliorate the effects of loss of *dip-2* function.

To determine whether lack of either DIP-2 or SAX-2 affected TAT-5 expression, we tagged the endogenous *tat-5* gene, which encodes multiple isoforms with two different N-termini. A GFP KI to TAT-5B and TAT-5D (*wur36*) showed broad expression in the nervous system and other tissues including oocytes (Fig S7B). We specifically examined TAT-5 expression in ALMs. TAT-5 puncta were present in the soma and along the ALM axon (Fig 4E). In *dip-2(0)* and *sax-2(0)* single mutants, the intensity of TAT-5 soma puncta was lower than in control, and some puncta were found in ectopic posterior neurites (Fig 4E,F). These results suggest that TAT-5 is re-located to ectopic neurites from the soma in the absence of DIP-2 or SAX-2.

We also examined whether PAD-1(E1909K) affected TAT-5 expression. In both neurons and oocytes, GFP::TAT-5B/D appeared largely unchanged by *pad-1(E1909K)* (Fig S7B,D), consistent with previous findings that in embryos, PAD-1 did not alter TAT-5 localization or vice versa ^29^. However, we noticed that in oocytes, PAD-1(E1909K) KI showed more co-localization with TAT-5B/D KI, compared to the wild type PAD-1 KI (Fig S7C). This finding suggests that PAD-1(E1909K) may preferentially promote TAT-5 activity at the membrane.

### *pad-1(E1909K)* suppresses *dip-2 sax-2* double mutant defects in plasma membrane morphology

We next investigated whether PAD-1(E1909K) suppressed morphological defects of neurons through regulation of plasma membrane structure. We visualized the TRN plasma membrane using a GFP::PH (PLC-δ1) marker (Fig 5A). ALM neurons in *dip-2(0)* single mutants displayed ectopic posterior neurites but otherwise normal GFP::PH distribution, whereas in *sax-2(0)* single mutants GFP::PH localized to abnormal vesicles within ALM soma or at the surface of neurons (arrows) (Fig 5A,B). *dip-2(0) sax-2(0)* double mutants displayed increased numbers of aberrant GFP::PH vesicles at the cell surface of TRNs (arrows). *pad-1(E1909K)* mutants displayed normal GFP::PH morphology as single mutants, and suppressed TRN membrane marker defects in *dip-2(0) sax-2(0)*. These results suggest that DIP-2 and SAX-2 function together to maintain neuronal plasma membrane morphology and that PAD-1(E1909K) can restore normal membrane morphology in the absence of *dip-2* and *sax-2*.

**Figure 5.**
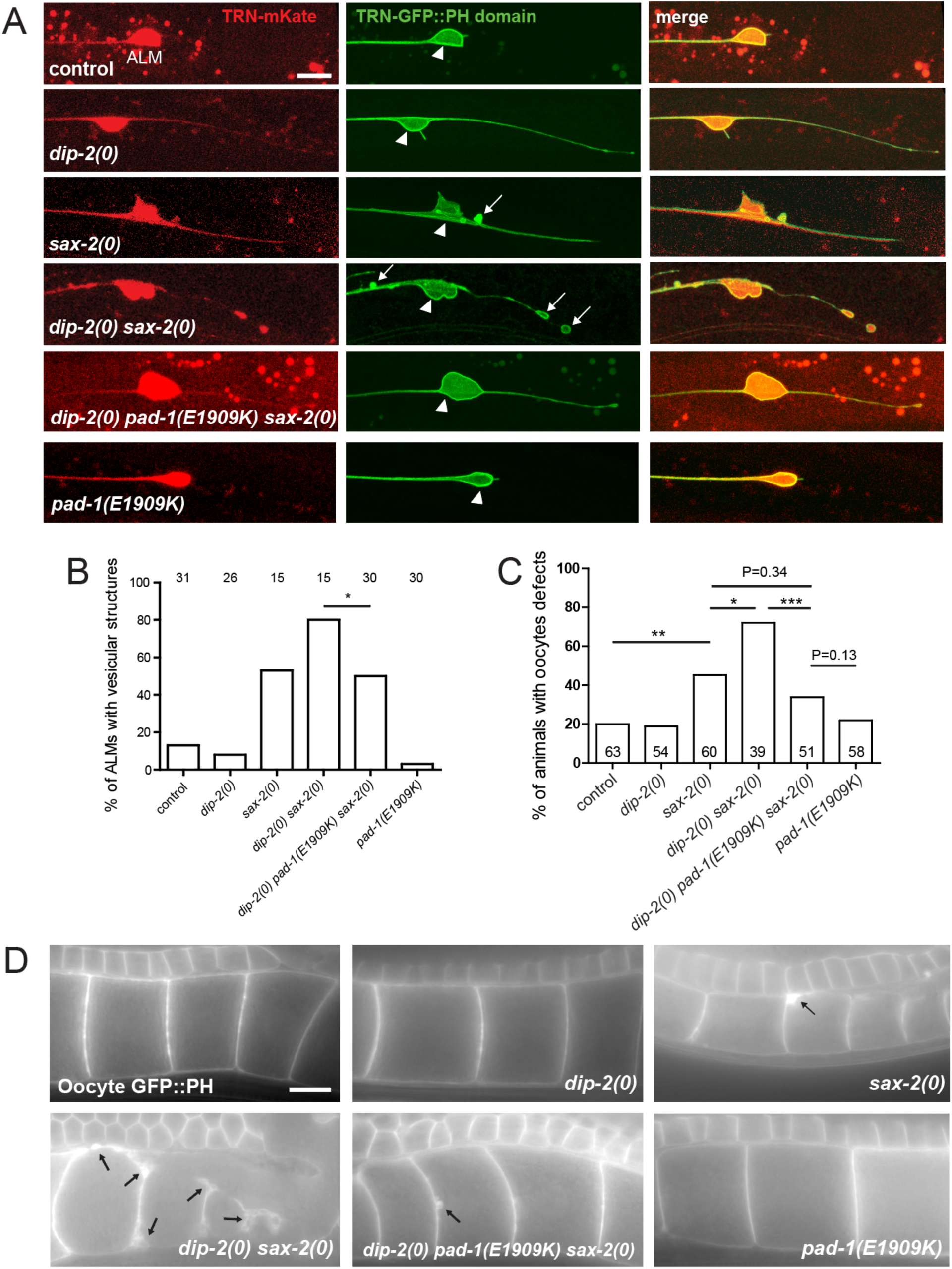
DIP-2 and SAX-2 synergistically regulate neuronal and oocyte membrane morphology. (A) Confocal images (maximum intensity projections of 5-10 focal planes) of ALM neurons labelled with mKate and GFP::PH domain under the control of the *mec-4* promoter (*juEx8228*). White arrows indicate vesicular structures labelled with GFP::PH; arrowheads indicate ALM soma. Scale = 10 µm. (B) Quantitation of ALMs exhibiting GFP::PH vesicular structures in the soma and proximal neurites (see Methods). Statistics: Fisher exact test. *(P<0.05) (C) Quantitation of oocyte membrane defects in 1-day old young adults. Statistics: Fisher’s exact test. (D) Images show oocyte plasma membranes labelled with *ltIs38* [P*pie-1*-GFP::PH]. *dip-2(0)* single mutants displayed normal membrane morphology. *sax-2(0)* single mutants and *dip-2(0) sax-2(0)* double mutants displayed variable abnormalities including thickened membrane patches and vesicles close to or outside the oocyte plasma membrane; these defects were suppressed to near wild type by *pad-1(E1909K),* which as a single mutant displayed normal plasma membrane morphology. Black arrows indicate aberrant vesicular structures or plasma membrane patches. Scale = 10 µm.

We further examined oocyte plasma membrane structure. In *dip-2(0)* single mutants, oocyte membrane morphology as visualized by a GFP::PH marker showed a thin and uniform distribution resembling that of wild type, whereas *sax-2(0)* single mutant oocytes displayed thickened patches of GFP::PH (Fig 5D). In *dip-2(0) sax-2(0)* double mutants, oocyte membrane morphology was severely disrupted with >70% of animals displaying thickened membrane patches or large vesicles outside oocytes; oocytes were also variable in size (Fig 5C,D). *pad-1(E1909K)* suppressed the oocyte membrane defects of *dip-2(0) sax-2(0)* double mutants and displayed normal oocyte membrane morphology as visualized by GFP::PH in single mutants. Thus, DIP-2 and SAX-2 act together to ensure plasma membrane morphology in oocytes as in neurons, and PAD-1(E1909K) can counteract the effects of loss of DIP-2 and SAX-2.

### *pad-1(E1909K)* and *tat-5(A316V)* suppress extracellular vesicle release in non-neuronal cells and neurons

PAD-1 and TAT-5 repress EV release from embryonic cells ^29^. We next tested whether PAD-1(E1909K) or TAT-5(A316V) affect EV release from the plasma membrane in embryos. Using a proteasomal degradation-based assay to visualize EVs labelled with a plasma membrane reporter ^35^, we found that *pad-1(E1909K)* or *tat-5(A316V)* single mutants exhibited normal embryonic EV release (Fig 6A). The GFP::PAD-1 N-terminal KI (*babIs1*) causes partial loss of function, resulting in abnormally elevated EV release in embryos ^32^. Introduction of E1909K into the GFP::PAD-1 background suppressed this elevated EV release to near wild-type levels (Fig 6A). These data show that the E1909K gain-of-function potentiates the ability of PAD-1 to inhibit EV release in embryos, suggesting the restoration of membrane morphology in *dip-2(0) sax-2(0)* double mutants could be due to reduced EV release.

**Figure 6.**
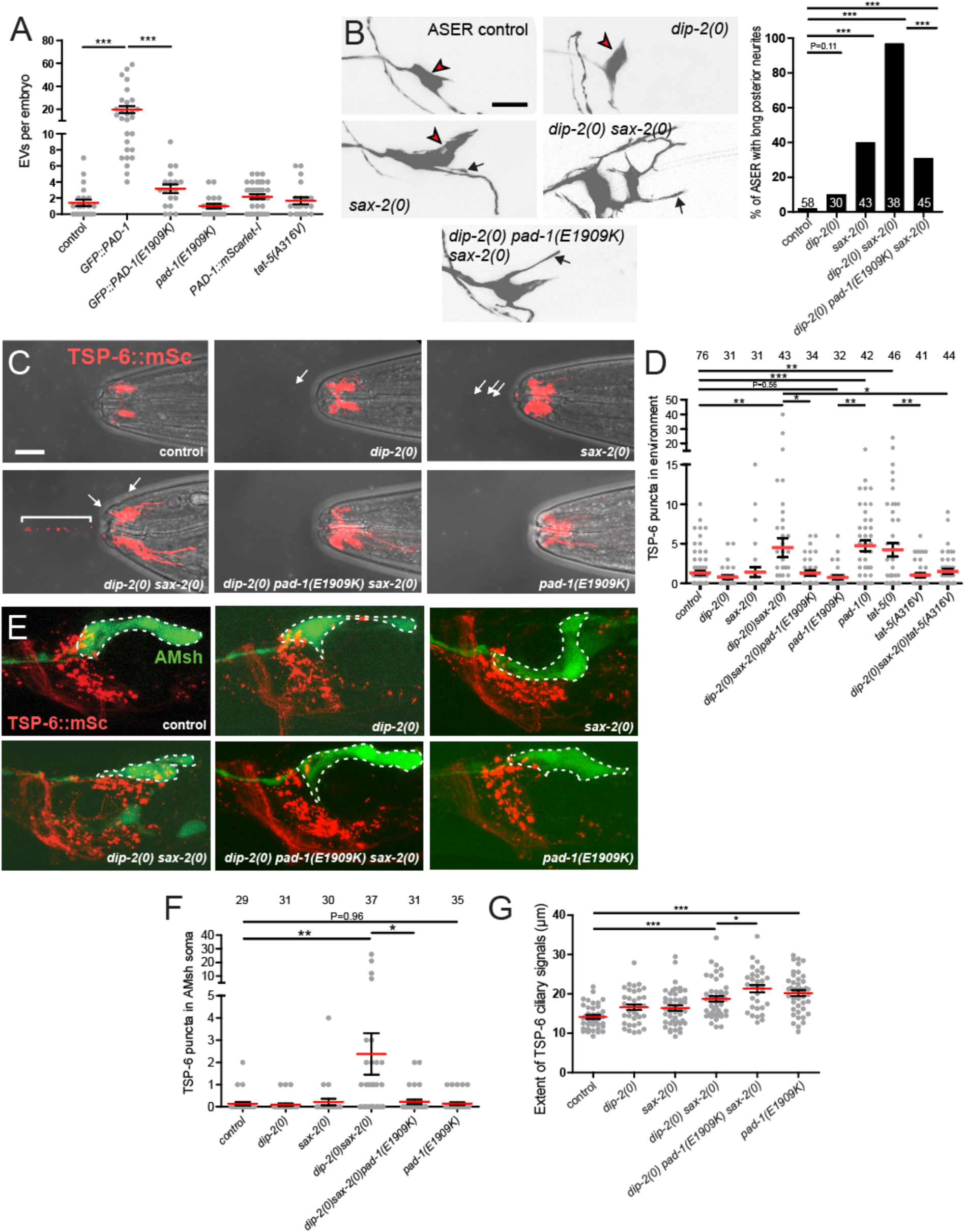
Elevation of TSP-6+ neuronal EVs in *dip-2(0) sax-2(0)* double mutants and suppression by *pad-1(E1909K)* and *tat-5(A316V)* (A) Quantitation of EVs released from embryonic cells. EVs labeled by mCherry::PH::CTPD were elevated in the partial loss-of-function GFP::PAD-1 KI; this was suppressed by introduction of *E1909K* mutation*. pad-1(E1909K)* and PAD-1::mSc KI displayed normal levels of embryonic EVs. Statistics, t-test with Bonferroni correction. (B) Images and quantitation of morphological defects in ASER neuron (marker *ntIs1*). In *sax-2(0)* single mutants, ASER displayed ectopic neurite outgrowth (black arrows) and misshapen soma shape (red arrowheads). *dip-2(0)* mutants displayed largely normal ASER morphology and strongly enhanced *sax-2(0)* phenotypes; the synergistic defects of *dip-2(0) sax-2(0)* double mutants were suppressed by *pad-1(E1909K)*. Statistics, Fisher’s exact test. (C) Confocal images of environmental EVs (TSP-6::mSc-+ puncta), defined as puncta released from the nose tip and observed in the adjacent medium. Superimposed confocal and bright field to show nose tip morphology. White arrows and bracket indicate TSP-6::mSc-+ EVs. (D) Quantitation of environmentally released TSP-6::mSc EVs. Statistics: one-way ANOVA, Šidák post test. (E) Images of the nerve ring region, head neurons expressing TSP-6::mSc, and AMsh cells labelled with YFP (outlined in dashed lines). TSP-6::mSc puncta were occasionally seen in the AMsh of *dip-2(0)* or *sax-2(0)* single mutants. In *dip-2(0) sax-2(0)* double mutants, TSP-6::mSc puncta accumulated in the soma of AMsh. *pad-1(E1909K)* suppressed the accumulation of TSP-6::mSc puncta in the AMsh soma and had normal levels of TSP-6::mSc puncta as single mutant. (F) Quantitation of TSP-6::mSc puncta in the soma of AMsh glial cells. Scale = 10 µm. Statistics: Kruskal-Wallis test, Dunn post test. (G) Quantitation of extent of TSP-6::mSc localization in ciliated neurons. Statistics: Kruskal-Wallis test with Dunn’s post test. For all the above statistical results, *** (P<0.001), ** (P<0.01), *(P<0.05).

To examine how PAD-1 affected EV release in neurons, we focused on ciliated sensory neurons, which generate several types of EVs ^36,37^ and which display aberrant morphology in *sax-2(0)* mutants ^11,12^. Morphology of the ciliated chemosensory neuron ASER was severely disrupted in *dip-2(0) sax-2(0)* double mutants compared to single mutants, and these morphological defects were significantly suppressed by *pad-1(E1909K)* (Fig 6B). Thus, the interactions between DIP-2, SAX-2, and PAD-1 observed in TRN morphology also apply to ciliated neuron morphology. To address their effects on EV production we examined the CD9-related tetraspanin TSP-6, which marks EVs released from ciliated neurons, which exit the *C. elegans* body through the amphid pore into the environment or are taken up by adjacent glial cells (amphid sheath cells, AMsh)^38^. Using a TSP-6::mSc KI marker, we found that *dip-2(0)* or *sax-2(0)* single mutants displayed normal levels of TSP-6+ environmental EVs released from ciliated neurons (Fig 6C,D). In contrast, *dip-2(0) sax-2(0)* double mutants displayed significantly elevated levels of environmental EVs (Fig 6C,D), as well as increased TSP-6+ puncta accumulation in the AMsh soma (Fig 6E,F). Loss of function in *pad-1(0)* or *tat-5(0)* resulted in elevated levels of TSP-6+ environmental EVs (Fig 6D). *pad-1(E1909K)* displayed normal levels of environmental EVs as a single mutant, but significantly suppressed both TSP-6+ EV release and TSP-6+ puncta accumulation in AMsh in *dip-2(0) sax-2(0)* double mutants (Fig 6C-F). Similarly, *tat-5(A316V)* single mutants exhibited normal levels of EVs and suppressed the enhanced release of TSP-6+ EVs in *dip-2(0) sax-2(0)* double mutants (Fig 6D). Together, these results suggest that loss of function in DIP-2 and SAX-2 results in increased release of neuronal EVs, and that these defects are suppressed by gain-of-function in PAD-1 or TAT-5.

While analyzing TSP-6::mSc expression in sensory neurons, we noticed that in *dip-2(0) sax-2(0)* double mutants, TSP-6::mSc was localized to a larger cellular surface area compared to control and each single mutant (Fig 6C,G). Interestingly, *pad-1(E1909K)* single mutants also exhibited an expanded zone of TSP-6+ membrane compared to controls (Fig 6C,G). This expansion of TSP-6+ localization in *pad-1(E1909K)* may reflect aberrant retention of TSP-6 in the neuronal plasma membrane. Collectively, these results show that PAD-1 and TAT-5 coordinately regulate EV release and neuronal morphology.

## Discussion

Maintenance of neuronal morphology throughout life requires a complex interplay of cytoskeletal and membrane dynamics. We found that two conserved proteins, DIP-2 and SAX-2, converge on a common mechanism that maintains the morphology of neurons and other cells. We identified novel gain-of-function mutations in the membrane trafficking regulator PAD-1 and the phospholipid flippase TAT-5, each of which can significantly improve neuronal morphology in the absence of DIP-2 and SAX-2. Our findings uncover a previously unknown network by which DIP-2 and SAX-2 regulate maintenance of membrane morphology via PAD-1 and TAT-5.

### DIP-2 and SAX-2 function synergistically in neurons and other tissues

Our genetic analysis reveals specific synergistic interactions between DIP-2 and SAX-2 in maintenance of neuronal morphology. The neuronal defects observed in *dip-2(0) sax-2(0)* double mutants are generally a combination of more severe or more penetrant versions of the single mutant phenotypes, suggesting that DIP-2 and SAX-2 play partly redundant roles in maintenance of neuronal morphology. The orthologs of DIP-2 function in lipid metabolism ^25^, whereas SAX-2/Fry family members are thought to have multiple functions including as scaffolds for SAX-1/NDR kinases ^20^. We find that overexpression of DIP-2 did not compensate for loss of SAX-2. These findings suggest that the two proteins are not redundant in biochemical function but converge on a common process.

DIP-2 and SAX-2 function synergistically in multiple tissues including the epidermis and the germ line. Moreover, gain-of-function of the PAD-1 and TAT-5 pathway suppresses *dip-2(0) sax-2(0)* synergistic defects in all tissues examined, supporting a common cellular mechanism. SAX-2/Fry family members have been implicated in morphogenesis of non-neuronal tissues in other organisms: Furry was first identified from its role in hair morphogenesis in *Drosophila* ^39^, whereas FRY functions in gastrulation in *Xenopus* ^40^ and in mammary gland development in mice ^41^. DIP2 family members may also function in multiple tissues, as mouse Dip2b is widely expressed including in epithelia ^42^. Thus, DIP2 and FRY family members could interact in multiple cell types in metazoa.

DIP-2::GFP localizes to the plasma membrane of epithelial cells and oocytes and is diffuse in the cytosol of neurons, whereas SAX-2::GFP forms intracellular puncta that do not co-localize with the reporters for several subcellular compartments. SAX-2::GFP puncta also do not co-localize with PAD-1::mSc in oocytes or neurons. Thus, SAX-2 could function in distinct subcellular compartments from DIP-2 and PAD-1. In *Drosophila*, overexpressed FRY-GFP forms motile puncta ^43^. We have not observed SAX-2::GFP puncta to be motile under our imaging conditions; however, the mechanisms localizing SAX-2 are an avenue for future investigation.

### Role of PAD-1 and the effect of E1909K

The strong genetic synergism of *dip-2* and *sax-2* mutations provided a sensitized background to screen for suppression of non-neuronal defects (shortened body length and reduced fertility) irrespective of effects on neuronal morphology. Among suppressors selected based on restoration of body length and fertility, we have analyzed *pad-1(E1909K)* and *tat-5(A316V)* in detail as these caused strong suppression of neuronal defects. Both single mutants are superficially wild-type and display essentially normal neuronal morphology, validating the approach of screening for suppression of non-neuronal phenotypes as a means to find rare mutations that may not be isolated based on their neuronal phenotypes.

PAD-1/Dopey proteins have multiple functions in membrane trafficking: in *C. elegans* embryos, PAD-1 inhibits the release of ectosome-type EVs from the plasma membrane and promotes phagocytosis of cell corpses and debris by regulating the activity of TAT-5 flippase ^29,32^. PAD-1 and TAT-5 are also thought to mediate endosomal trafficking as enlarged late endosomes are observed in the corresponding mutants ^29^. Supporting this previous finding, we show that PAD-1 in neurons partially co-localizes with RAB-7 in the endosomal trafficking pathway (Fig 3B). As the E1909K mutation increases PAD-1 plasma membrane localization in some cell types, it may primarily affects the plasma membrane function of PAD-1.

Dopey family proteins are large proteins (1700-2400 aa) with multiple conserved domains or regions. The highly conserved N-terminal ‘Dopey’ domain is involved in interaction with kinesin light chain ^30^, whereas the central region is implicated in interactions with other proteins such as Mon2 or flippases ^44^. The C-terminus appears to be involved in membrane association ^30^. In mammalian Dopey1, a ‘Dopey Extreme C-terminus’ (DEC) region was sufficient for PI4P binding ^30^, yet the equivalent region of Dopey2 was insufficient for PI4P binding. PAD-1 is approximately equally similar in sequence to mammalian Dopey1 and Dopey2, and it remains to be determined whether its localization mechanism resembles either mammalian ortholog. The E1909 residue lies outside of these previously studied domains but in a well-conserved region predicted to form a series of alpha helices. Our data indicate that this region of PAD-1 is essential for function and that the E1909 residue affects plasma membrane localization. The E1909K or E1909A changes might alter the activity of this domain via lipid binding, Dopey dimerization or self-inhibition, or interaction with a factor in the plasma membrane, such as TAT-5.

### The function of TAT-5 and the effect of A316V

The flippase TAT-5 was initially characterized in *C. elegans* embryos ^33^. Loss of function of TAT-5 leads to externalized PE, as shown by increased staining with the PE binding lantibiotic Duramycin. *tat-5(0)* mutant embryos are lethal and show increased plasma membrane budding, reduced phagocytosis, disrupted cell shape and motility, indicating that PE asymmetry is likely critical for normal membrane morphology of the embryo. Interestingly, loss of function in *pad-1* results in higher PE exposure than loss of function in *tat-5*, suggesting PAD-1 regulates TAT-5 and at least one other flippase or scramblase ^29^. PE asymmetry has not yet been analyzed in *C. elegans* neurons due to a lack of genetically encoded PE sensors. During postembryonic development, TAT-5 and PAD-1 also function in Q cell migration via retromer-dependent sorting of Wntless^45^. As such, TAT-5 and PAD-1 may play a role in intracellular membrane trafficking in addition to regulation of EV release.

*tat-5(A316V)* suppresses *dip-2(0) sax-2(0)* phenotypes. As a single mutant, *tat-5(A316V)* animals are viable and fertile, and have overtly normal neuron morphology in young and day 5 adults, whereas in *tat-5(0)* mutants ALM neurons show long ectopic posterior neurites by day 5 of adulthood (Fig 4C,D). These differences suggest A316V does not reduce TAT-5 function. As TAT-5 is widely expressed in the nervous system, it is possible that TAT-5(A316V) may act cell-autonomously in neurons. A316 lies in the actuator (A) domain and is conserved in the essential subclass of P4-ATPases, with the equivalent residue A375 in *S. cerevisiae* Neo1p on the surface of the A-domain ^34^. As the A domain is involved in the dephosphorylation step of the ATPase cycle, A316V might alter phosphatase activity, potentially enhancing the flippase activity of TAT-5. Protein structural and biochemical experiments will be required to dissect mechanistically how A316V alters TAT-5 function.

### Neuronal membrane morphology, lipid asymmetry and EV biogenesis

Our results suggest that DIP-2 and SAX-2 function convergently to regulate lipid composition or membrane trafficking. As yet it is unclear if *dip-2(0) sax-2(0)* double mutants have overall altered lipid composition or a more specific defect such as altered plasma membrane lipid asymmetry. Plasma membranes are highly asymmetric in lipid distribution ^46^, and phospholipid asymmetry has been implicated in maintenance of neuron morphology in several contexts. Externalized phosphatidylserine (PS) is widely known to act as an ‘eat-me’ signal to cause glial pruning in mammalian synapses ^47^, and can also act as a ‘save-me’ signal to promote fusion of axon fragments after injury^48^. In *C. elegans,* externalized PS can sculpt sensory neurites ^49^. Loss of function in the major PS flippase TAT-1 causes PS exposure and results in loss of TRNs via increased phagocytosis of entire TRNs ^50^ and enhanced neurite engulfment in AFD sensory neurons ^49^. Although we have not comprehensively investigated the relationship of the DIP-2 or SAX-2 pathway to PS-mediated pruning, some observations suggest that these two are mechanistically distinct. For example, TRNs in *dip-2(0) sax-2(0)* double mutants are present throughout life. Moreover, loss of function in the phagocytic receptor CED-1 does not result in aberrant TRN morphology ^51^, nor does compound loss of function of Ca^2+^-dependent lipid scramblases ANOH-1/2 that disrupt PS asymmetry ^52^. Speculatively, the defects in *dip-2* and/or *sax-2* mutants might reflect altered PE asymmetry.

The mechanism by which DIP-2 and SAX-2 convergently regulate membrane trafficking and EV release remains to be elucidated. One possibility is that DIP-2 and SAX-2 each act on multiple membrane flippases. Alternatively, the combined loss of DIP-2 and SAX-2 could alter the level of lipids that activate TAT-5, as many flippases are activated by their cargo lipids or specific non-cargo lipids ^34,53–55^. In either case, an increase in externalized PE could result in excessive membrane curvature, and consequently elevated levels of EV production. The functional relationship of membrane lipid asymmetry and EV release in neuronal morphology is of great interest. In ciliated neurons, EV release maintains normal morphology by removing deleterious excess of ciliary material ^38^, reminiscent of the proposed role of much larger ‘exopher’-type EVs in maintaining TRN proteostasis and mitostasis ^56^. In contrast, our data suggest that elevated EV release in *dip-2(0) sax-2(0)* double mutants correlates with aberrant neuronal morphology, and that suppression of neuronal morphology and membrane defects by *pad-1(E1909K)* correlates with suppression of EV release to normal levels. EV release may promote or impair neuronal morphology depending on the level or type of EVs involved. Alternatively, and not mutually exclusive, EVs from defective neurons might play roles in communication with other neurons or tissues. Notably, our findings suggest a common mechanism for regulation of EV production in neuronal and non-neuronal cell types.

The genes studied in this work are highly conserved across animals, and human orthologs have been implicated as candidates for neurological disorders. DIP2A and DIP2B have been linked to neurocognitive disorders ^24,57^, and FRY has been linked to intellectual disability and developmental delay ^58^. DOPEY2 is located on human chromosome 21 and has been investigated for its possible contribution to Down Syndrome ^59^, whereas DOPEY1 loss of function in rats causes defective myelin formation ^60^. Loss of function in the TAT-5 ortholog ATP9A results in aberrant neurite morphology and endosomal recycling defects in mice ^61^, and ATP9A variants have been linked to human neurodevelopmental disorders ^62,63^. Our findings taken together with results from other model organisms suggest that a DIP2/FRY/DOPEY/Flippase pathway could be involved in maintenance of neuronal morphology in humans, and that altered activity of this pathway may contribute to neurological disorders. Further genetic analysis of *dip-2 sax-2* signaling in *C. elegans* could identify additional relevant factors in this pathway.

## Methods

### Key Resource Table

| Reagent or resource | Source | Identifier |
| --- | --- | --- |
| Bacterial strains |  |  |
| *Escherichia coli* OP50 | *Caenorhabditis* Genetics Center | RRID: WBStrain00041969 |
| Experimental models: organisms/strains |  |  |
| *C. elegans*, see Table S1 | This paper | N/A |
| Oligonucleotides |  |  |
| Primers for genotyping and sequencing and cloning, see Table S2 | This paper | N/A |
| crRNAs for gene editing, see Table S3 | This paper | N/A |
| Recombinant DNA |  |  |
| pID3.01 | Addgene | 22720 |
| pDD104 | Addgene | 47551 |
| pCFJ2412 | Jorgensen lab | N/A |
| pTG95 | Min Han lab | N/A |
| Molecular Biology Reagents |  |  |
| TOPO-TA cloning kit | Thermo Fisher | K250020 |
| LR Clonase II mix | Invitrogen | 12538-120 |
| Gibson Assembly Master Mix | NEB | E2611 |
| Imaging |  |  |
| Levamisole | Sigma | L9756 |
| Agar | BD | 214040 |
| Agarose | Fisher | NC0493156 |
| Software |  |  |
| Fiji (ImageJ2) | Open Source | https://imagej.net/software/fiji/ |
| Slidebook 6 | 3i | [intelligent-imaging.com](https://www.intelligent-imaging.com) |
| Prism 9 | GraphPad | [graphpad.com](https://www.graphpad.com) |

### Experimental Model

Wild-type *C. elegans* used in this study is N2 Bristol ^64^. Strains were maintained at 20°C on Nematode Growth Media (NGM) plates seeded with *E. coli* OP50. Strains were constructed using standard procedures including crossing using males, visually detecting behavioral or morphological phenotypes, and genotyping by PCR, superselective PCR ^65^, or sequencing for mutant alleles.

### Method details

*C. elegans* strain genotypes, primers, newly generated plasmids, CRISPR reagents, and allele sequence information are listed in in Supplemental Tables 1-5. Genetic strains were constructed by standard crosses and alleles verified by PCR or sequencing.

### Transgene construction and germ line transformation

To generate a TRN plasma membrane marker P*mec-4*-GFP::PH, the rat PLC PH domain was amplified from *ltIs38* [P*pie-1*-GFP::PH ^66^] using primers AC4641 and AC4642 (Supplemental Table 2) and subcloned into pCR8 vector by TOPO-TA cloning. LR reaction was performed between the resulting pCR8 vector and pCZGY603 [P*mec-4*-GFP] ^67^. The final construct pCZGY3591 (Supplemental Table 3) was injected into wild type N2 at concentrations ranging from 1 ng/µl to 20 ng/µl, with P*ttx-3*-dsRed (10 ng/µl) as a co-injection marker and pBluescript (50 ng/µl) as filler DNA. After titrating concentrations of the construct, 2 ng/µl was determined as the optimal concentration to generate multiple transgenic animals displaying normal morphology of TRNs. *juEx8228* was crossed to mutant strains to examine TRN membrane morphology.

We generated transgenes that express sense and antisense RNAs ^68^ to specifically knock down *pad-1* in TRNs. A 1.4 kb fragment of *pad-1* cDNA corresponding to exons 8-10 was amplified using AC4633 and AC4634. The sense promoter for *mec-4* gene was amplified using AC4637 and AC4639; the anti-sense promoter was amplified using AC4637 and AC4640. The *pad-1* cDNA fragment was mixed with the sense promoter and fusion-PCRed using AC4638 and AC4636. The fragment was also mixed with the antisense promoter and fusion-PCRed using AC4638 and AC4635. The two fusion-PCR products were injected at 10 ng/µl into *pad-1(ju1806) zdIs5; sax-2(ju1815)* with P*myo-2*-mCherry as co-injection marker, generating *juEx8255* and *juEx8256*.

### CRISPR gene editing

For deletions and point mutations we generally used a *dpy-10* co-CRISPR strategy ^69^. For knock-in gene editing we used a self-excising cassette strategy ^70^.

#### sax-2

The *sax-2(ju1815)* deletion was generated using crRNAs targeting exon 2 and the penultimate exon (see Supplemental Table 4). The resulting deletion conferred phenotypes similar to *sax-2(ky216)* in neuronal morphology (Fig S1C); *dip-2(0) sax-2(ju1815)* was not significantly different in body length from *dip-2(0) sax-2(ky216)* (Fig S2B). The *sax-2(ju1815)* deletion was fully recessive. A GFP::SAX-2 N-terminal knock-in was obtained from SunyBiotech (*syb3389*). The homology-directed repair (HDR) plasmid for C-terminal SAX-2::GFP(*ju1831*) was made using three-fragment Gibson Assembly (New England Biolabs) in which two *sax-2* homology arms containing 498 bp upstream of the *sax-2* TAA stop codon and 540 bp downstream of the *sax-2* TAA stop codon plus the stop codon were amplified from N2 genomic DNA and fused such that they flanked GFP (pDD282). Sequencing of the insertion revealed that homologous recombination had introduced a premature stop codon in the loxP site 5’ to the FLAG tag, generating 132 bp between this stop codon and 3’ UTR. The premature stop codon might potentially affect expression of the *ju1831* KI. The SAX-2 KIs did not display overt phenotypes as single mutants and did not affect body length in *dip-2(0)* backgrounds.

#### pad-1

To edit E1909K(*ju1881, ju1882, ju1987)*, E1909A(*ju1933*), or the 8-residue deletion (*ju1948*) in *pad-1*, a mix of *pad-1* crRNA, *dpy-10* crRNA, tracerRNA, Cas9 protein, and a *pad-1* repair oligo was injected into strains of interest: the two KI strains GFP::PAD-1 (MCP6) and PAD-1::mScarlet-I (WEH655) and CZ26881. Dpy F1s were singled and screened for the presence of E1909K mutation using superselective primers (Supplemental Table 2). F2s with the homozygous mutation were outcrossed twice then if necessary balanced with *tmC27* ^71^. The PAD-1::mScarlet-I C-terminal knock-in (*syb4112*) was obtained from SunyBiotech; mScarlet-I was Germ Line Optimized (GLO) ^72,73^, with a linker QCFFIMPTLYKKAGCS between *pad-1* and mScarlet-I. The PAD-1::mSc KI strain displayed normal embryonic EV levels (Fig 6A). The PAD-1::mNeonGreen C-terminal knock-in (*syb6942*) was obtained from SunyBiotech; an auxin-inducible degron (AID) was inserted between PAD-1’s C-terminal end and mNeonGreen.

#### tat-5

A mix of *tat-5* crRNA, *dpy-10* crRNA, tracerRNA, Cas9 protein, *tat-5* repair oligo containing the same mutation as in *ju1805* (GCA to GTA at codon 316 in isoform A) was injected into a control strain (CZ10175, *zdIs5*) and into the *dip-2(0) sax-2(0) zdIs5* double mutant strain (CZ27791). F1 Dpy animals were singled and screened for the presence of a Pvu II restriction site introduced via the repair oligo. F2 animals with the homozygous mutation were obtained and outcrossed twice. The resulting edits were named *ju1877* and *ju1878* respectively. The GFP::TAT-5B/D KI *wur35* was generated by NemaMetrix in an *unc-119(ed3)* mutant strain. *C. elegans* optimized GFP with *rpl-18* intron 2 and 150 bp synthetic PATC introns from pCFJ2412 ^74^ with a floxed *unc-119* selection cassette in an intron and a flexible linker from pID3.01 (Addgene 22720) were inserted into the upstream operonic SL2 ATG of *tat-5b* and *tat-5d* using pNU2343. pDD104 (Addgene 47551) and pTG95 (P*sur-5*-NLS-GFP) were injected to remove the selection cassette with Cre recombinase and generate *wur36*, which was then outcrossed twice to N2.

### Genetic screen for new *dip-2-*like mutants altering TRN morphology

To identify new mutations affecting maintenance of TRN morphology, we screened 3-day old adult animals of genotype *fem-3*(*hc17*ts); *zdIs5*[P*mec-4*-GFP] adults for aberrant *dip-2-*like ectopic neurites. Following standard 50 mM EMS mutagenesis, F1 progeny were transferred to new NGM plates (3 F1/plate) and allowed to lay eggs at 20°C. Each individual F2 generation plate consisting of L3 and L4 stage animals was split, with half allowed to grow at 20°C and half shifted to the restrictive temperature of 25°C. Animals at 25°C were assessed for neuronal morphology 3 to 4 days later when most were 3-day old adults. Single animals were recovered from the corresponding plate maintained at 20°C, propagated for several days and then rescreened using epifluorescence on a Zeiss SteREO Discovery V8. We screened ∼7200 haploid genomes and identified 11 mutants (Fig S1A; Supplemental Table 5). New *dip-2*-like mutants were outcrossed at least 3 times and genomic DNA prepared for whole genome sequencing (WGS) as described ^75^. WGS was performed at the Centre for Applied Genomics, Toronto. In the course of these studies, we found that several “*eno*” mutants isolated on the basis of ectopic neurite outgrowth in motor and sensory neurons ^76^ were allelic to *dip-2* and phenotypically resemble previously reported *dip-2(0)* alleles (Supplemental Table 5).

### Genetic screen for suppressors of *dip-2 sax-2* synergistic defects

We mutagenized L4 animals of strain CZ27791 *dip-2(gk913988) zdIs5; sax-2(ky216)* with 50 mM EMS following standard protocols ^64^. We screened F2 progeny for normal body morphology (‘non-Dpy’) as well as improved growth and reproduction. We classified candidate suppressors by TRN soma morphology: isolates with abnormal TRN soma were classified as ‘*dip-2* specific’ and isolates with restored TRN soma morphology were classified as ‘*sax-2* specific’. Some suppressors were difficult to assess for the specificity of their neuronal suppression. The two strongest candidate suppressors were outcrossed to N2 or *sax-2(ky216)* and mapped.

Whole genome sequencing (BGI Americas) was obtained for CZ27791, the starting un-mutagenized strain; CZ27891, the outcrossed *dip-2 sax-2* double mutants suppressed by *ju1806*; and CZ28020, the original isolate of *dip-2 sax-2* double mutants suppressed by *ju1805* (Supplemental Table 1). The Galaxy platform was used to analyze raw sequence files with a custom-designed workflow ^77^; homozygous variants in the three strains were determined by comparison to WormBase reference N2 sequence (ce10) and to a reference lab copy of N2 (CZ21293); variants present in CZ21293 were not considered further. Variants with quality scores above a threshold of 100 and a coverage of 10 were kept for further analysis.

In mapping *ju1805* and *ju1806,* variants present in CZ28020 or CZ27891 but not in CZ27791 were first extracted from the whole genome sequencing result. In the meantime, unsuppressed or suppressed recombinants were re-isolated after outcrossing with *sax-2(ky216)*; suppressed recombinants were readily re-isolated in the F2 generation and unsuppressed recombinants were re-isolated in the F2 and F3 generations from several independent outcrosses. Analysis of recombinant lines indicated *ju1805* and *ju1806* were linked to *dip-2* on chromosome I. Among SNPs in this region we confirmed that *ju1806* changes *pad-1* codon 1909 (GAA to AAA) to convert glutamic acid to lysine and *ju1805* changes *tat-5* codon 316 (GCA to GTA) to convert alanine to valine.

### Quantitation of body length, brood size, and oocyte morphology

For measurement of body length, images of animals were taken using a Zeiss Axioplan Imager compound microscope and imported into ImageJ (Fiji). Segmented lines were drawn longitudinally from the mouth to the posterior end of intestine in animals such that these lines were at equal distance from dorsal and ventral sides of animals at any point of the lines. Measured lengths were normalized to appropriate controls. For brood size assay, one L4 animal was placed on a plate, allowed to lay eggs for a day, and transferred to a new plate every day for four consecutive days, and the number of eggs laid in each of five plates was counted and combined to obtain the brood size. A Zeiss Axio Imager M2 compound scope was used to score oocyte morphology in day 1 adults. Oocyte plasma membranes labelled with *ltIs38* ^66^ were examined for thickening, folding, or presence of internal or external vesicles.

### Microscopy

For imaging of neuronal morphology and worm body, animals were immobilized in M9 buffer with 2 mM levamisole and mounted on a 4% agar pad. All confocal fluorescence images were acquired with a 63x/NA1.4 Plan-Aprochromat objective lens using Zeiss LSM800 confocal microscope. For green fluorophores, 488 nm laser power was set at 5% and detector gain at 500-800 V. For red fluorophores, 561 nm laser power was set at 22% and detector gain at 600-800 V. z stacks were acquired with 3-12 slices set 0.5-1 µm apart. Maximum intensity projections were performed using Fiji ImageJ. Bright field images showing body length and morphology were taken either on a Zeiss Axio Imager M2 compound microscope or Zeiss LSM800 confocal microscope at 10x magnification under identical settings.

For Zeiss AiryScan imaging, animals were immobilized with 10 mM levamisole and mounted on a 10% agarose pad. Images were acquired using a Plan-Apochromat 63x/NA 1.40 oil immersion objective using Zeiss LSM900 microscope (Axio Observer.Z1/7) equipped with Airyscan 2. In Zen 3.4(Blue) software, SuperResolution mode with ‘Best Signal’ detection setup was selected. The laser for green fluorescence was set at 488 nm (470-525) for wavelength, 1.5-5% laser power and 800 V detector gain. The laser for red fluorescence was set at 561 nm (550-617), 12-15% power, and 800 V detector gain. Scans were in frame mode with unidirectional scan direction and scan speed of 7. For Airyscan SR imaging mode we used autofilter with Super Resolution values ranging from 7.5-8.4.

### Characterization of neuronal morphology

A Zeiss Axio Imager M2 compound microscope was used for visual scoring of defects in neuronal morphology of live animals. Animals were anesthetized using 5 mM levamisole in M9 buffer. Markers used are: P*mec-4*-GFP(*zdIs5*), P*mec-7*-GFP(*muIs32*), and P*unc-25*-GFP(*juIs76*). TRN morphology was scored using *zdIs5* unless otherwise noted. In scoring ALM ectopic neurite outgrowth, any projections from the ALM soma other than the anterior axon were measured and scored as “ectopic posterior neurite” if longer than 10 µm. ‘Multiple ectopic neurites’ was defined as >1 ectopic neurite extending from the ALM soma. PLM overshooting was defined as the axon of PLM extending anterior to the ALM soma ^13^. ALM soma shape was scored as normal (0), mildly defective (1) or severely defective (2) as detailed in Figure 2 legend.

### Quantitation of PAD-1 fluorescence intensity in oocytes and nerve ring

Images of oocytes expressing GFP::PAD-1, PAD-1::mScarlet-I (mSc), or PAD-1::mNeonGreen (mNG) were acquired using LSM800 confocal and imported into Fiji (ImageJ). A straight line was drawn from the center of nucleus to that of an adjacent oocyte so that the midpoint of the line is positioned at the interface of plasma membranes of the two oocytes. The intensity along the line was measured in Plot Profile and the peak intensity value (i.e. plasma membrane) was averaged. For quantitation of intensity signal at granules/recycling endosomes, the diameter of a granule was drawn, and the average intensity value of the diameter was measured and then further averaged. 5 granules were randomly selected per oocyte and quantified. For quantitation of cytosol intensity signal, three lines, each of which is 2-5 µm in length, were drawn without crossing any granules in the cytoplasm of each oocyte and their average intensity determined. For quantitation of PAD-1::mNG and GFP::TAT-5B/D in nerve ring, a box was drawn in the neuropil between the pharynx and the ventral nerve cord. The mean fluorescence intensity in the box was measured and normalized to control.

### Imaging and quantitation of neuronal TSP-6::mSc-containing extracellular vesicles

To image neuronal EV release we used TSP-6::mScarlet (*syb4122*) ^38^. Animals were immobilized with 10 mM levamisole in M9 buffer and mounted on a 4% agarose pad for 1 h to achieve complete paralysis. Animals of multiple genotypes were mounted on the same slide to reduce variation due to imaging conditions. Confocal fluorescence images were acquired in a z-stack with a 63x objective lens on a Zeiss LSM800 confocal microscope. Maximum intensity projections were performed using Fiji. For quantitation of environmentally released EVs, we manually counted puncta labelled with TSP-6::mSc in a 67.25 µm (x) x 39.91 µm (y) x 12-17 µm (z) z-stack adjacent to the mouth. For quantitation of EVs accumulated in the soma of AMsh cells, we manually counted any red puncta found in the soma throughout the z-stack. Under our imaging conditions TSP-6::mSc EV production was similar to that previously reported ^38^. To quantify the extent of TSP-6::mSc neuronal localization, a straight line was drawn longitudinally from the most anterior signal to the most posterior signal in each animal, and the length of this line was measured.

### Quantitation of embryonic extracellular vesicles

Individual EV puncta between the cell surface and eggshell were counted from images of live 3- to 15-cell embryos expressing the PH::CTPD marker ^35^ using Slidebook6 (3i). Thick clusters or patches of EVs were counted as 1-2 puncta depending on size and EV puncta were not counted between cells, likely undercounting the number of released EVs.

### Statistical analysis

Statistical analyses were performed using GraphPad Prism 9. Statistical significance was determined using Fisher’s exact test, unpaired t test, one-way ANOVA or Kruskal-Wallis tests followed by a post-hoc test. Data are represented as mean ± SEM unless noted. Sample sizes are shown inside or above columns in figures or legends.

## RESOURCE AVAILABILITY

### Lead contact

Further information and requests for resources and reagents should be directed to and will be fulfilled by the Lead Contact, Andrew Chisholm.

### Materials Availability

All unique/stable reagents generated in this study are available from the Lead Contact without restriction.

### Data Availability

Raw quantitative data will be made available in a source data file. Representative image files will be deposited in the Figshare repository.

## Acknowledgements

We thank members of Jin and Chisholm labs for valuable inputs throughout the work, Yue Sun and Junxiang Zhou for advice on whole genome sequencing and mapping. We thank Junxiang Zhou for the AMAN-2 marker, Zhiping Wang for TRN specific RAB-5 and RAB-7 transgenes, Ken Noma for the P*rgef-1*-mCh transgene, Erin Jyo, Alyssa Zhang, Charlotte Sue, Katharina Beer, and Patrick Flores for strain construction. We thank Patrick Laurent for TSP-6 reagents and advice, Michael Ailion and Erik Jorgensen for neuronal Golgi and endosomal markers. Some strains were provided by the CGC, which is funded by NIH Office of Research Infrastructure Programs (P40 OD010440). S.P was a trainee on the UCSD Neural Circuits Training Grant T32 NS007220 (P.I., N. Spitzer). Funding: NIH R01 NS093588 to A.D.C and Y.J., Natural Resources and Engineering Research Council of Canada grant 2018-06790 to A.C. and N.N., Deutsche Forschungsgemeinschaft (DFG) grant WE5719/2-1 to A.M.W., and a grant from the Paul G. Allen Family Foundation to A.M.W. and A.D.C.

## Supplemental Figures

**Supplemental Figure 1.**
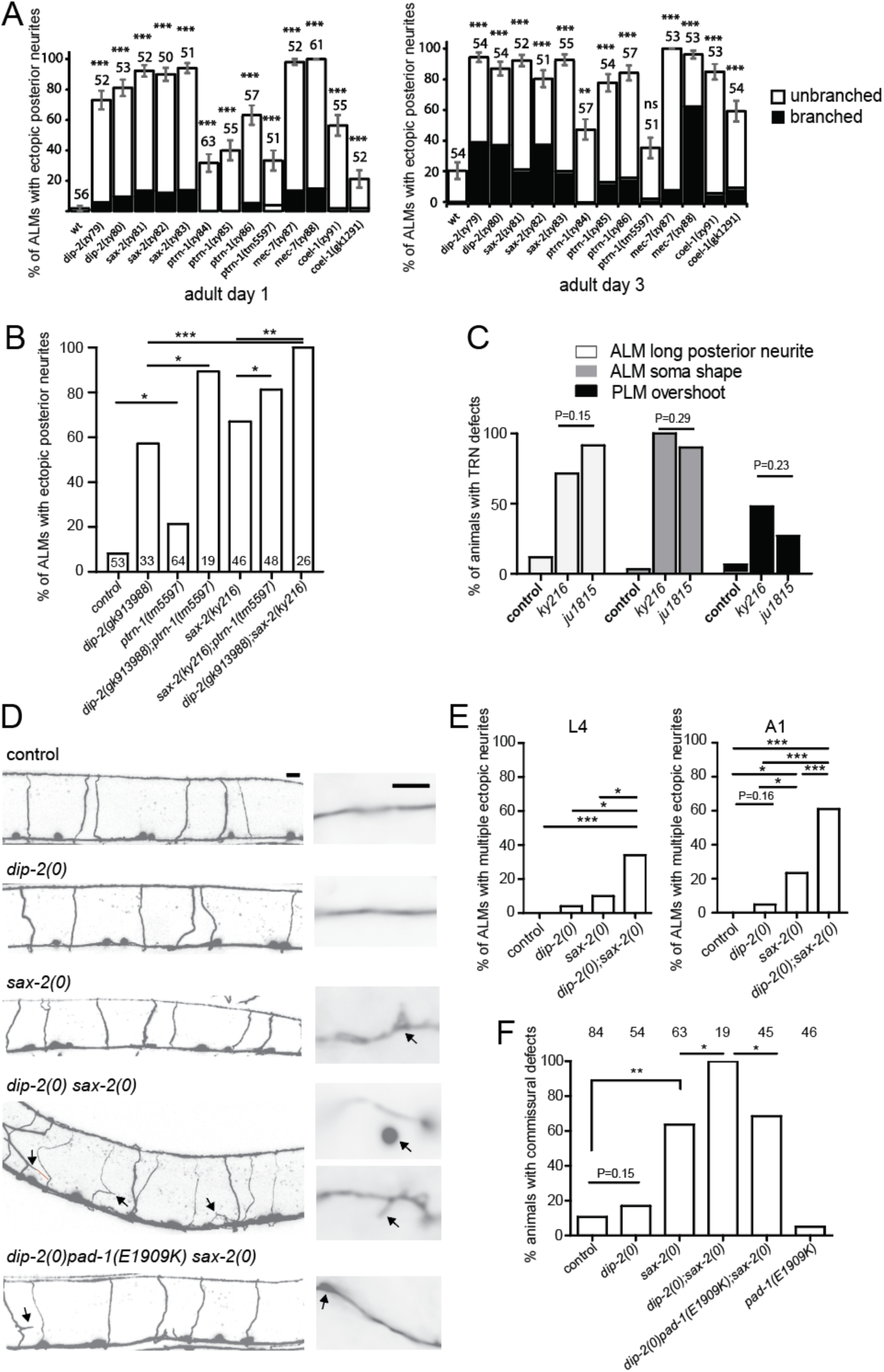
d*i*p*-2* displays specific synergistic interaction with *sax-2* in maintenance of neuron morphology. (A) Quantitation of ALM ectopic posterior neurites in mutants at 1-day (*left*) and 3-day (*right*) of adulthood. Mutants with *dip-2-*like phenotypes were isolated in a forward mutagenesis screen. Ectopic neurites were classified as branched or unbranched. Statistics for A: bars show mean and error bars indicate SEs of proportion for the indicated number of animals. Significance is compared to controls (*zdIs5*) using a one-way ANOVA with Tukey test for multiple comparisons. (B) Genetic interactions between *dip-2* and null mutations in genes involved in TRN neuronal morphology maintenance. *dip-2* and *sax-2* displayed the strongest synergistic interaction. ALM ectopic neurites were scored in 1-day old adults, 1 ALM per animal, transgenic marker *zdIs5*. N, number of ALMs scored. (C) Comparison of between *sax-2(ju1815)* deletion and *sax-2(ky216)* in TRN defects. Both mutations caused similar levels of morphological defects. N = 40-50 per genotype. (D) Confocal images (*left*) of D-type GABAergic motor neurons labelled with P*unc-25*-GFP(*juIs76*). *dip-2(0)* single mutants did not exhibit significant defects in D-type neuron morphology; in *sax-2(0)* single mutants, D-type neurons occasionally exhibited kinks, blebs, or round protrusions in their lateral commissures. In *dip-2(0) sax-2(0)* double mutants, D-type motor neurons displayed fully penetrant defects in commissural morphology as well as ectopic neurite sprouting in the ventral nerve cord (arrows). Scale = 10 µm. Details of commissure morphology are shown in enlarged insets (right); scale = 1 µm. Black arrows indicate defects including ectopic neurites, blebs, and vesicular structures. Confocal z-stacks were projected with maximum intensity. (E) Quantitation of ALM neurites with multiple branches in animals of genotype indicated. *dip-2(0) sax-2(0)* double mutants displayed synergistic increases in multiply branched ALM neurites in L4 and day 1 adult (A1). (F) Quantitation of commissural defects as % of animals with at least one defective commissure, scored in L4 stage. Statistics for panels B,C,E, and F: Fisher’s exact test. *** (P<0.001), ** (P<0.01),* (P<0.05).

**Supplemental Figure 2.**
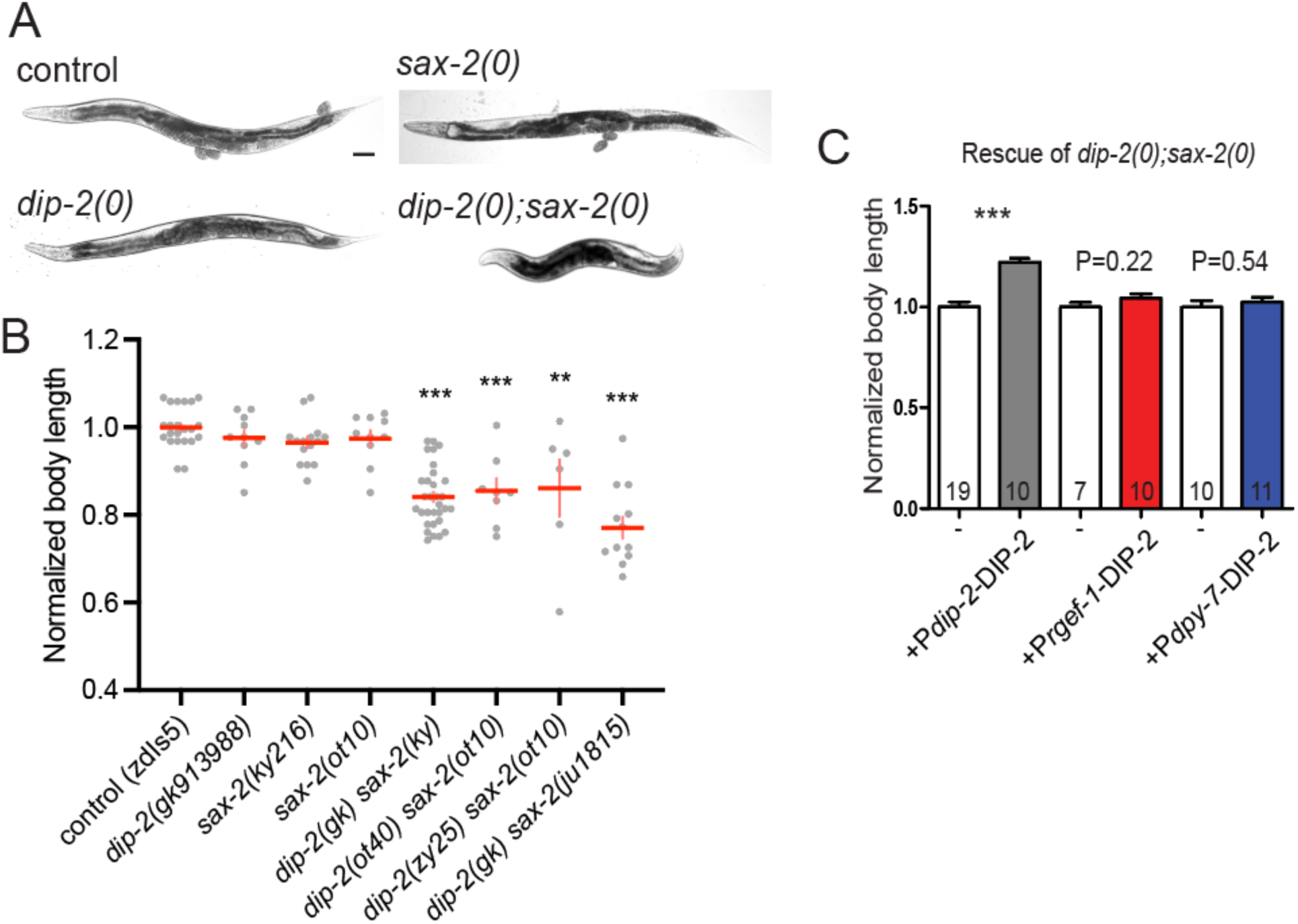
d*i*p*-2 sax-2* double mutants display synergistic effects on body length. (A) Bright field images of 1-day old adults of genotype indicated. *dip-2(0)* or *sax-2(0)* single mutants have normal morphology and body length, whereas *dip-2(0) sax-2(0)* double mutants are shorter in body length. Scale = 100 µm. (B) Quantitation of normalized body length in 1-day old adult animals; *zdIs5* background except for *dip-2(gk913988) sax-2(ju1815)*. All *dip-2 sax-2* double mutant combinations were significantly different from controls and were not significantly different from each other. N = 6-31 per genotype. Statistics: one-way ANOVA with Tukey post test. *** (P<0.001). (C) Quantitation of body length in 1-day old adult *dip-2(0) sax-2(0)* double mutants with and without transgenes, normalized to *dip-2(0) sax-2(0)* control. Body length defects of *dip-2(0) sax-2(0)* double mutants were rescued by overexpression of DIP-2 using its own promoter but not using the pan-neuronal *rgef-1* or the epidermal *dpy-7* promoters. Statistics: t test. *** (P<0.001).

**Supplemental Figure 3.**
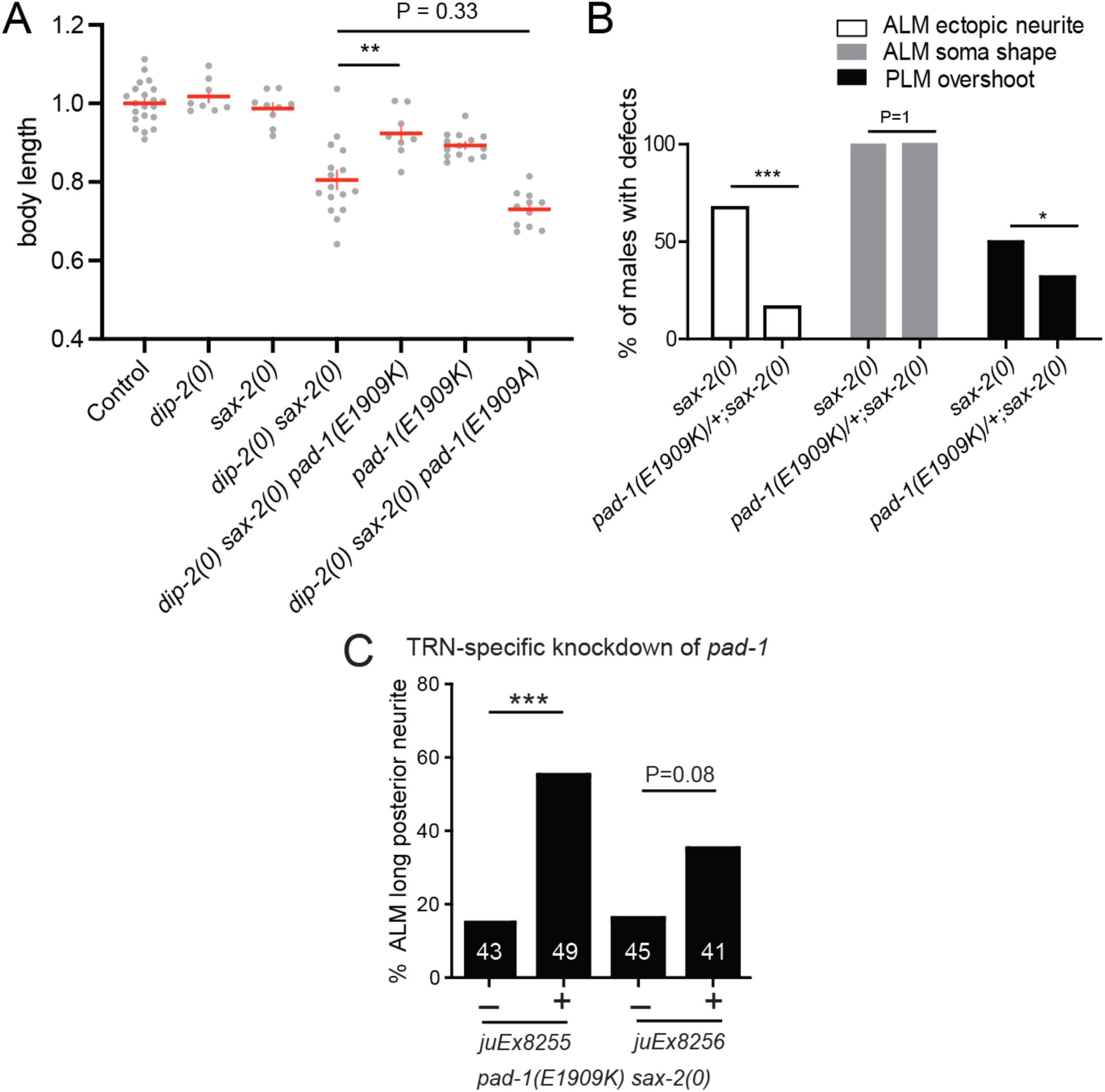
Suppression of *dip-2 sax-2* double mutant and *sax-2* single mutant phenotypes by *pad-1(E1909K)* (A) Quantitation of body length in 1-day old adults. *pad-1(E1909K)* but not *pad-1(E1909A)* suppressed the short body length of *dip-2(0) sax-2(0)* double mutants. N = 5-15 per genotype; normalized to wild type controls. Statistics: t-test. (B) Quantitation of morphological defects of touch neurons in *sax-2(0)* mutant males with heterozygous *pad-1(ju1806)*. *pad-1(ju1806)* acted as a semi-dominant suppressor of *sax-2* ALM ectopic neurite and PLM overshooting defects. N= 45-59. Statistics: Fisher exact test. *** (P<0.001),* (P<0.05). (C) TRN-specific dsRNA-mediated knockdown (KD) of *pad-1* reverses the suppression of ectopic neurite outgrowth. *pad-1* KD in *pad-1(ju1806) sax-2(0)* increased % of ALMs with ectopic neurite outgrowth. Statistics: Fisher exact test. ***(P < 0.001), ** (P<0.01),* (P<0.05).

**Supplemental Figure 4.**
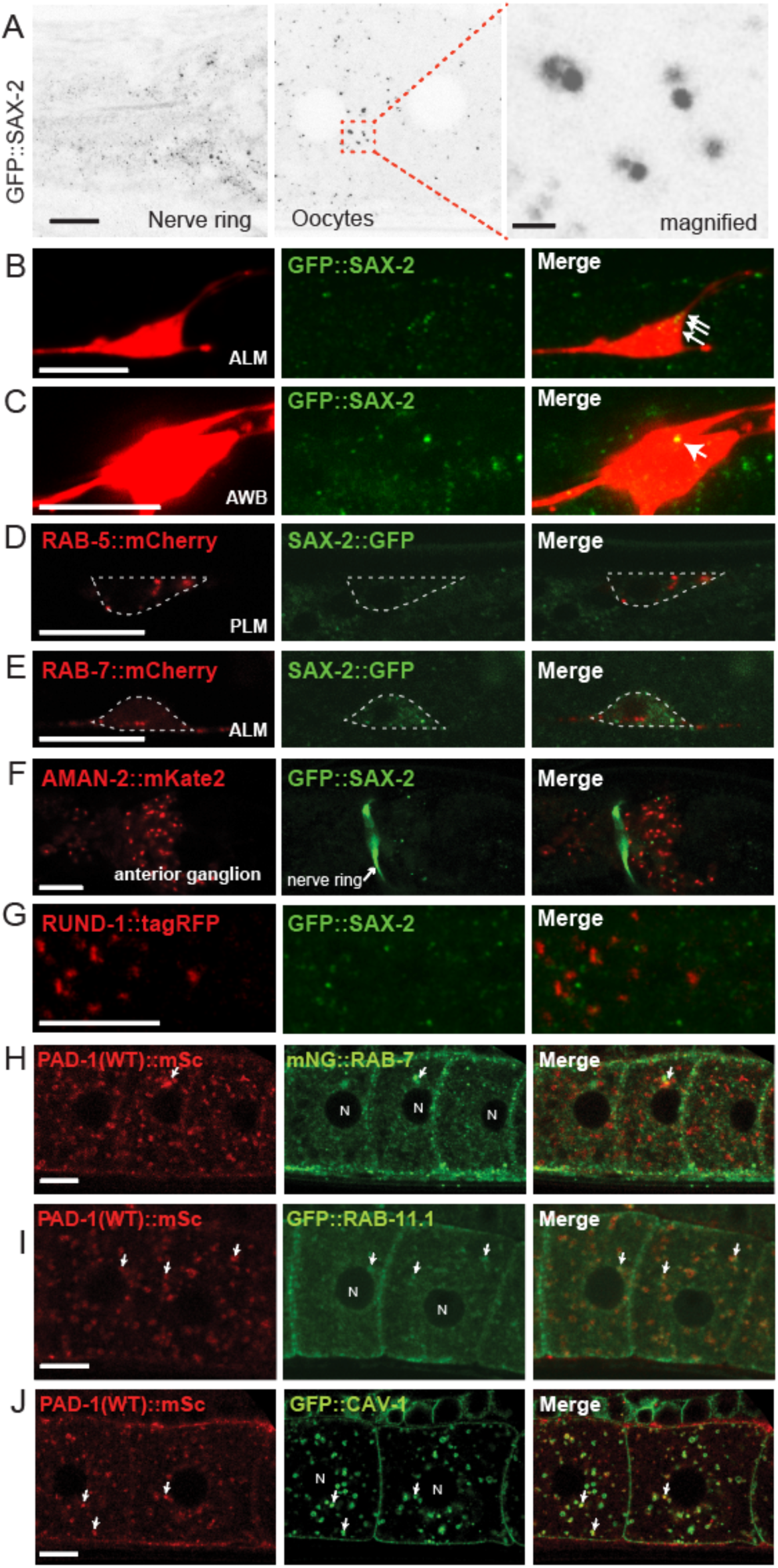
Localization of SAX-2::GFP to subcellular puncta in neurons and oocytes and lack of co-localization with endosome or Golgi markers. (A) Confocal images of GFP::SAX-2 (*syb3389*) knock-in expression in nerve ring and oocytes; maximum intensity projections of 3 focal planes, scale = 10 µm. Inset shows SAX-2 puncta magnified; scale = 1 µm. (B) GFP::SAX-2 forms small puncta (arrows) in peripheral soma of ALM neuron labeled with mCherry (*juEx5179*). (C) GFP::SAX-2 formed a single large punctum in soma of sensory neuron AWB (arrow; red marker = *oyIs65*). Maximum intensity projections of 15 focal planes, scale = 10 µm. (D-E) Confocal images of TRNs and nerve ring region co-expressing GFP::SAX-2 or SAX-2::GFP KI with red fluorescent markers: RAB-5::mCherry for early endosomes, RAB-7::mCherry for late endosomes, AMAN-2::mKate2 for medial Golgi, and RUND-1::tagRFP for *trans* Golgi. In images displaying the nerve ring labelled with GFP::SAX-2 and AMAN-2::mKate2 (*juEx8137*), AIY is labeled green due to P*ttx-3*-GFP co-injection marker. (H-J) Confocal images of oocytes co-expressing PAD-1::mSc and green fluorescent labels for intracellular organelles. PAD-1::mSc (white arrows) was partly co-localized with mNG::RAB-7 KI (*utx12*) and RAB-11.1 marking late and recycling endosomes, respectively. Vesicular PAD-1::mSc also partially co-localized with *pie-1-*driven GFP::CAV-1 (*pwIs28*), a component of caveolae and of cortical granules in oocytes ^79^. White arrows indicate punctate colocalization; N indicates nucleus. For panels B-J, scale = 10 µm.

**Supplemental Figure 5.**
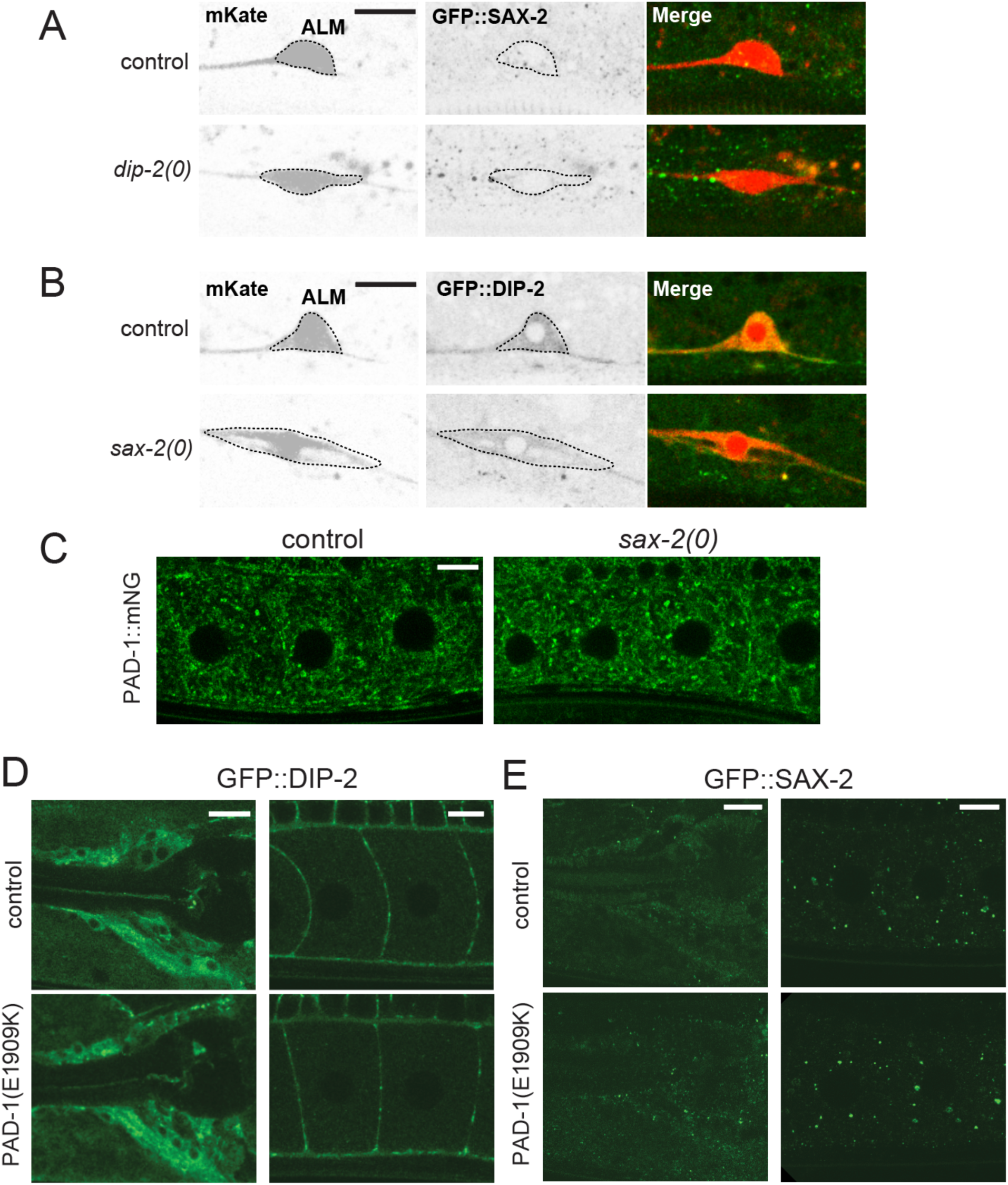
Localization of DIP-2, SAX-2, and PAD-1 in neurons and oocytes. (A) Confocal images of ALMs expressing GFP::SAX-2 KI. SAX-2 localization in ALM was not altered in *dip-2(0)* mutants. (B) GFP::DIP-2 KI localization in ALM was not altered in *sax-2(0)* mutants. (C) Confocal images of oocytes expressing PAD-1::mNG KI. In *sax-2(0)* mutants, PAD-1::mNG KI expression was not significantly different from wild type. (D) The expression pattern and level of GFP::DIP-2 in both neurons and oocytes were not altered in *pad-1(E1909K)* mutants. (E) PAD-1(E1909K) did not affect GFP::SAX-2 in both neurons and oocytes. For all panels, scale = 10 µm.

**Supplemental Figure 6.**
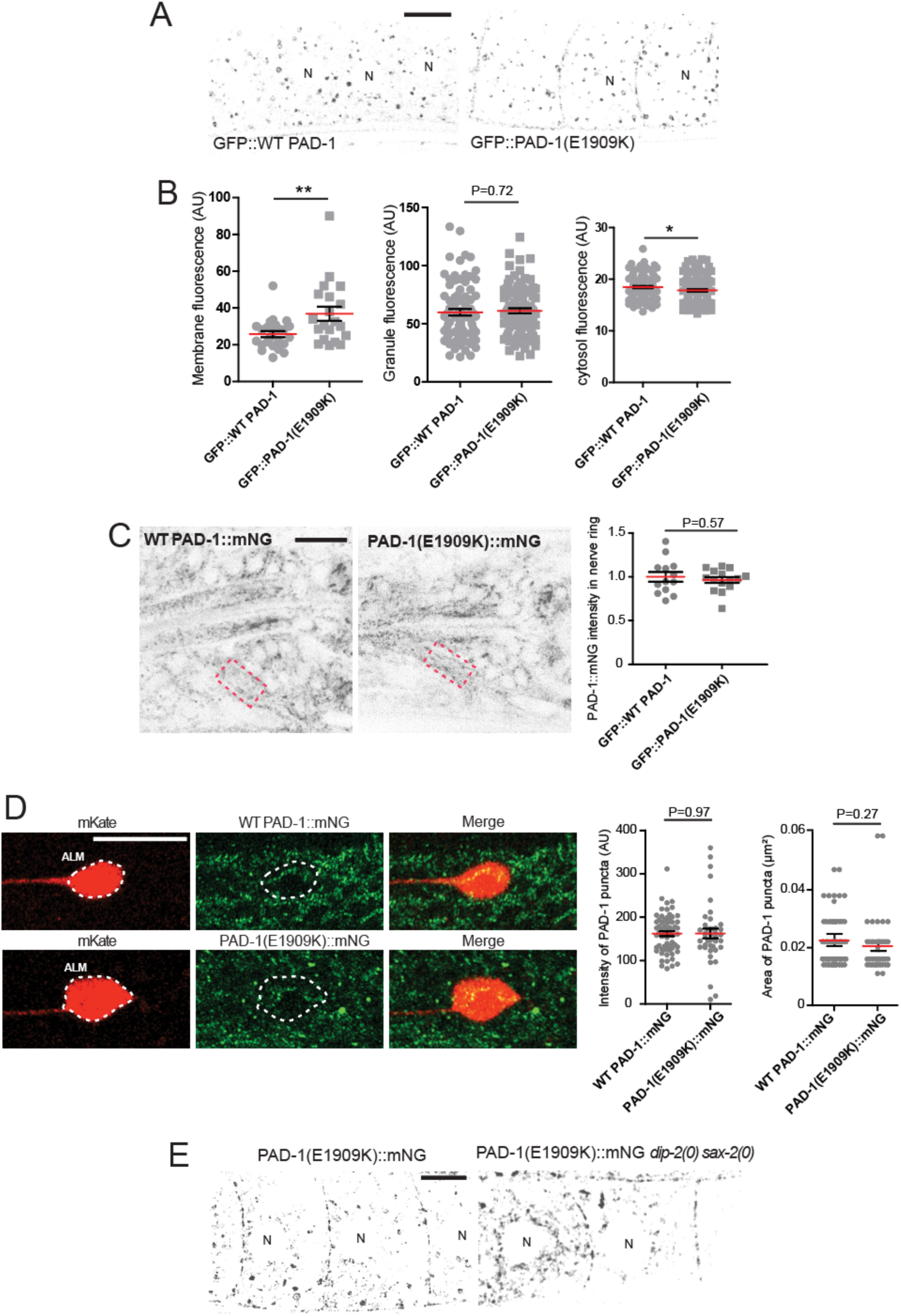
E1909K gain-of-function mutation augments PAD-1’s association with the plasma membrane in oocytes and the nervous system. (A,B) Confocal images and quantitation of GFP::PAD-1 localization and intensity. GFP::PAD-1 fluorescence at oocyte plasma membranes was significantly increased by E1909K; localization to vesicles/granules was unaffected and localization to cytosol was decreased. Dot plots show mean (red bar) and SEM (black). Statistics: t-test. ** (P<0.01),* (P<0.05). (C) Confocal images (lateral views) of anterior ganglia and nerve ring (red dashed boxes) and quantitation of normalized PAD-1::mNG expression levels in the boxed regions. E1909K did not increase PAD-1 nerve ring localization (t test). (D) Confocal images of ALM neurons labelled with mKate and PAD-1::mNG KI (WT and E1909K). Quantitation of PAD-1::mNG puncta intensity (arbitrary units, AU) and puncta size in ALM soma. Localization of PAD-1::mNG in ALM was not altered by E1909K. (E) Confocal images of PAD-1(E1909K)::mNG in oocytes where PAD-1 puncta were found in the cytoplasm and plasma membrane. The patchy pattern of PAD-1(E1909K)::mNG puncta in oocyte plasma membrane was not altered in *dip-2(0) sax-2(0)* mutant background. For panels (A,C,D, and E), scale = 10 µm.

**Supplemental Figure 7.**
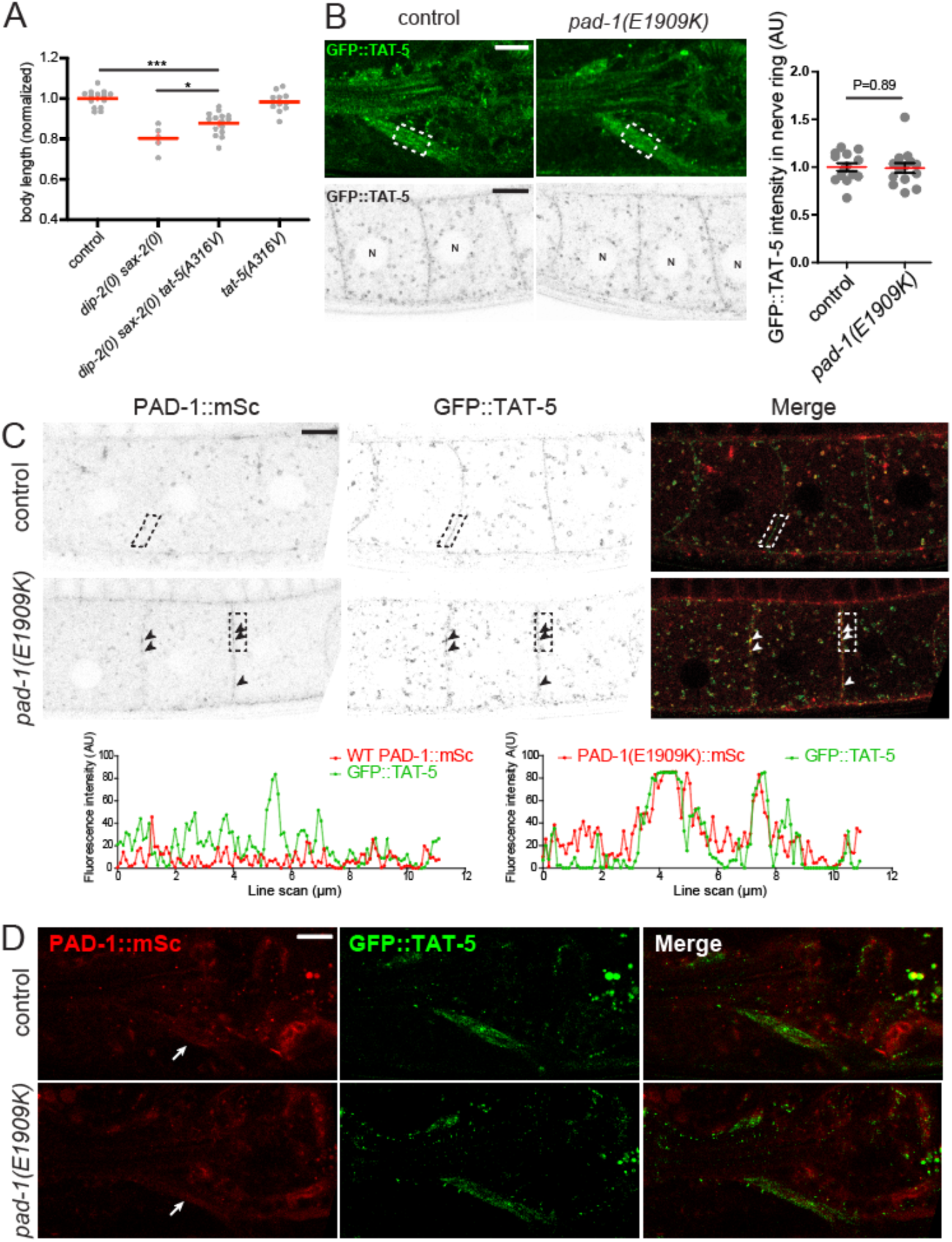
Suppression of *dip-2 sax-2* phenotypes by *tat-5(A316V)* and TAT-5 localization in *pad-1(E1909K)* (A) Quantitation of normalized body length measured in 1-day old adults. *tat-5(ju1878 A316V)* partially suppressed the reduced body length of *dip-2(0) sax-2(0)* double mutants and had normal body length as single mutant. Statistics: one-way ANOVA with Tukey’s post test. *** (P<0.001),* (P<0.05). (B) Confocal images of GFP::TAT-5B/D KI (*wur36*) in the nervous system (*top*) and oocytes (*bottom*) and quantitation of GFP::TAT-5B/D in nerve ring (*right*). GFP::TAT-5 localized to axons in the nerve ring (dashed boxes). Dot plot, normalized fluorescence signals of GFP::TAT-5 in the nerve ring. GFP::TAT-5 nerve ring localization was normal in *pad-1(E1909K)* mutants (t test). In oocytes, GFP::TAT-5 localized to punctate vesicles and close to the plasma membrane; this localization was similar in *pad-1(E1909K).* Oocyte nuclei indicated by N. (C) PAD-1(E1909K)::mSc displayed increased co-localization with GFP::TAT-5 at the plasma membrane (confocal images; black arrow heads; white arrow heads in merge); both PAD-1(E1909K)::mSc and GFP::TAT-5 displayed patchy localization at the oocyte plasma membrane. Line scans of fluorescence at the plasma membrane within dashed boxes were plotted using Fiji to show increased co-localization of PAD-1(E1909K)::mSc with GFP::TAT-5. (D) GFP::TAT-5 localization in the nerve ring area was normal in PAD-1(E1909K)::mSc animals. White arrows indicate nerve ring. For all panels, scale = 10 µm.

